# Motor Neuron Dysfunction in SORD Deficiency: Implications for Therapeutic Development in Peripheral Neuropathies

**DOI:** 10.64898/2026.05.13.724849

**Authors:** Giuseppina Divisato, Stefano Tozza, Emanuela Cascone, Elena Polishchuk, Maria Chiara Zizolfi, Emilia Giannino, Fiorella Marsella, Daniela Di Girolamo, Ciro Menale, Lucia Perone, Paolo Gianfico, Giovanni Cuda, Cecilia Bucci, Paolo Maiuri, Roman Polishchuk, Fiore Manganelli, Silvia Parisi

## Abstract

Biallelic mutations in the sorbitol dehydrogenase (*SORD*) gene have been identified as one of the most common causes of autosomal-recessive Charcot–Marie–Tooth disease type 2 (CMT2) and distal hereditary neuropathy, collectively referred to as SORD deficiency. These mutations result in loss of sorbitol dehydrogenase activity, a key enzyme in the polyol pathway that metabolizes glucose, leading to marked accumulation of sorbitol in patient-derived fibroblasts. However, the mechanisms by which SORD dysfunction drives axonal degeneration remain poorly understood, and robust in vitro models of human SORD-deficient motor neurons (MNs) are still lacking.

To address this gap, we established a human *in vitro* model of SORD deficiency by generating induced pluripotent stem cells (iPSCs) from fibroblasts affected individual carrying biallelic *SORD* mutations (*SORD*^c.757delG/c.316_425+165del^), and unaffected heterozygous carriers (*SORD*^c.757delG/wt^ and *SORD*^wt/c.316_425+165del^). These iPSCs were subsequently differentiated into motor neuron progenitors (MNPs) and MNs.

Comprehensive analysis of SORD-deficient human cells—including fibroblasts, MNPs, and MNs—revealed pronounced structural and functional abnormalities in the mitochondrial compartment, characterized by mitochondrial fragmentation and increased proton leak. Importantly, fibroblasts derived from two additional unrelated patients carrying the SORD mutation (*SORD*^c.757delG/c.757delG^) further confirmed that SORD deficiency is associated with a mitochondrial phenotype.

At the molecular level, SORD deficiency led to upregulation of aldose reductase (AR), another key enzyme of the polyol pathway, resulting in disruption of cellular redox homeostasis and increased oxidative stress. Consistent with these alterations, MNs derived from CMT2/SORD patients exhibited clear neurodegenerative features, including severe defects in neurite branching and synaptic architecture, ultimately impairing neuronal connectivity.

Notably, pharmacological inhibition of AR effectively rescued both mitochondrial dysfunction and neuronal structural defects, supporting the targeting of AR as a promising therapeutic strategy for polyol pathway-associated neuropathies as CMT2/SORD and diabetic neuropathy.

## Introduction

Charcot–Marie–Tooth (CMT) disease comprises a group of inherited peripheral neuropathies with a historical estimated prevalence of approximately 1 in 2,500 individuals^1^. Disease onset typically occurs during adolescence or early adulthood and is characterized by marked clinical and genetic heterogeneity. Common clinical features include *pes cavus*, progressive distal muscle weakness and atrophy, sensory loss, reduced or absent deep tendon reflexes, and gait impairment. Disability accumulates gradually over time, largely driven by progressive axonal degeneration. Based on electrophysiological and pathological features, CMT is broadly classified into demyelinating (CMT1) and axonal (CMT2) forms^2,3^. Distal hereditary motor neuropathy (dHMN) represents a subtype of CMT2 in which motor neurons are predominantly or exclusively affected^4,5^.

Despite relatively limited clinical variability, CMT exhibits extensive genetic heterogeneity, complicating both molecular diagnosis and the development of targeted therapies. To date, more than 130 genes have been implicated in CMT^6^. Among these, biallelic mutations in the sorbitol dehydrogenase (*SORD*) gene have recently emerged as a frequent cause of autosomal-recessive CMT2 (CMT2/SORD)^7^. Affected individuals typically present with phenotypes consistent with CMT2 or dHMN. Notably, the frameshift mutation *SORD*^c.757delG^ (p.Ala253GlnfsTer27), either in homozygous or compound heterozygous form, represents the most prevalent pathogenic variant associated with this condition^7^. Accordingly, genetic screening of *SORD* is now recommended in unresolved cases of axonal neuropathy, particularly CMT2 and dHMN^8,9^.

The molecular mechanisms linking *SORD* mutations to motor neuron degeneration remain incompletely understood. Fibroblasts derived from patients carrying biallelic *SORD* mutations exhibit markedly reduced SORD protein levels and accumulation of intracellular sorbitol, reflecting impaired conversion of sorbitol to fructose within the polyol pathway^7^. This pathway consists of two key enzymatic steps: the reduction of glucose to sorbitol catalysed by aldose reductase (AR), followed by oxidation of sorbitol to fructose mediated by sorbitol dehydrogenase^10^. A prevailing hypothesis is that intracellular sorbitol accumulation disrupts neuronal structure and function. Dysregulation of sorbitol metabolism has been extensively implicated in diabetic neuropathy, where hyperglycaemia-driven activation of the polyol pathway results in sorbitol overload^11^. This can perturb cellular redox homeostasis by depleting glutathione-related pathways and/or increasing the production of reactive oxygen species (ROS), ultimately promoting apoptosis and neuronal damage^12,13^. These mechanisms are thought to contribute to early neuronal dysfunction and structural impairment^14^. Consistently, patients with CMT2/SORD display elevated sorbitol levels in serum, urine, and fibroblasts^7,15^, while *Sord*−/− animal models develop motor-predominant neuropathy accompanied by increased sorbitol accumulation in serum, cerebrospinal fluid, and peripheral nerves^16^. Importantly, treatment with aldose reductase inhibitors (ARi), including epalrestat, ranirestat, and govarestat, reduces sorbitol levels and ameliorates neuronal phenotypes in experimental models^7,15^.

Nevertheless, the precise mechanisms by which SORD deficiency leads to axonal degeneration remain unclear. In the absence of SORD activity, sorbitol accumulation may recapitulate pathogenic processes observed under hyperglycaemic conditions or activate distinct disease-specific pathways. Notably, several genes implicated in CMT are involved in mitochondrial dynamics and function, highlighting mitochondrial dysfunction as a central feature of disease pathogenesis^17,18^. Consistent with this, studies in *Sord*-deficient *Drosophila melanogaster* suggest that sorbitol accumulation may impair mitochondrial integrity and function^15^, potentially contributing to degeneration of post-mitotic neurons.

To elucidate the mechanisms by which SORD deficiency induces MN degeneration, we generated and characterized a human *in vitro* model of CMT2 using induced pluripotent stem cell (iPSC)-derived MNs harboring different *SORD* genotypes (*SORD*^c.757delG/c.316_425+165del^, *SORD*^c.757delG/wt^, and *SORD*^wt/c.316_425+165del^). We demonstrate that SORD deficiency, as well as sorbitol accumulation, disrupts the mitochondrial network and activates apoptotic pathways. Furthermore, MNs carrying biallelic *SORD* mutations exhibit pronounced defects in neurite branching and synaptic connectivity. Notably, pharmacological inhibition of AR significantly rescues mitochondrial function and restores neuronal structural integrity.

Overall, our study identifies mitochondrial dysfunction and sorbitol accumulation as key drivers of SORD-related neuropathy and provides a robust human cellular model for mechanistic investigation and preclinical therapeutic development.

## Materials and Methods

### Patients

#### Patient#1

Firstborn of two siblings, born at term to healthy-looking non-consanguineous parents, after an uncomplicated delivery; psychomotor developmental milestones were normal. Since childhood, he has reported muscle pain and cramps predominantly affecting the lower limbs. During adolescence, he developed a tendency to trip frequently. At age 17, he sustained a fall resulting in a subtotal tear of the anterior talofibular ligament (ATFL). Since then, his gait has progressively worsened, with the onset of bilateral foot drop and progressive distal leg atrophy. The neurophysiological test showed an axonal sensory-motor neuropathy, namely CMT2.

At age 25, he presented with a steppage gait and was unable to walk on heels or toes. Examination showed mild hypotrophy and weakness of the first dorsal interosseous muscles, hypotrophy of the distal third of the legs, movements of the anterolateral compartment muscles possible against gravity, and mild weakness against resistance of the posterior leg muscles. Deep tendon reflexes were reduced in the upper limbs and absent in the lower limbs. Mild pes cavus and Achilles tendon retraction were noted. Pinprick sensation at the toes was reduced. The overall disability, assessed using the Charcot-Marie-Tooth Examination Score (CMTES)^19^, was mild (5/28).

#### Patient#2

Firstborn of two siblings, born at term to healthy non-consanguineous parents after an uncomplicated delivery; psychomotor developmental milestones were normal. She remained asymptomatic until her 50s, when she began to experience some difficulty in walking. Electrophysiological studies showed features of a degenerative axonal motor neuropathy, consistent with distal hereditary motor neuropathy (dHMN).

At age 55, she has a mild steppage gait, unable to walk on heels. Strength and deep tendon reflexes were normal in the upper limbs. In the lower limbs, examination showed hypotrophy of the distal third of the legs, with mild weakness of the tibialis anterior (MRC 4/5). Patellar reflexes were normal, while Achilles reflexes were absent. Mild pes cavus and Achilles tendon retraction were noted. Sensation (light touch, pinprick, vibration) was normal. The overall disability, assessed using the Charcot-Marie-Tooth Examination Score (CMTES), was mild (3/28).

#### Patient#3

The second of two siblings, born at term to healthy-looking non-consanguineous parents, after an uncomplicated delivery; psychomotor developmental milestones were normal. Onset of symptoms occurred in her 40s with difficulty walking. Electrophysiological studies showed features of a degenerative axonal motor neuropathy, consistent with distal hereditary motor neuropathy (dHMN). At age 51, she was unable to walk on tiptoes and had difficulty walking on heels. Strength and deep tendon reflexes were normal in the upper limbs. In the lower limbs, examination showed hypotrophy of the distal third of the legs, with mild weakness of the tibialis anterior (MRC 4/5) and marked weakness of the gastrocnemius (MRC 3/5). Patellar reflexes were normal, while Achilles reflexes were absent. Mild pes cavus and Achilles tendon retraction were noted. Sensation (light touch, pinprick, vibration) was normal. The overall disability, assessed using the Charcot-Marie-Tooth Examination Score (CMTES), was mild (3/28).

### Informed patient consent

Informed consent to participate in this study was obtained from three patients (referred to as SORD^-/-^ #1, SORD^-/-^ #2 and SORD^-/-^ #3 throughout the manuscript) and their respective family members (the father and mother of patient #1, referred to as SORD^+/-^ #1 and SORD^+/-^ #2 respectively throughout the manuscript) and a healthy, unrelated control (referred to as WT throughout the manuscript). Written informed consent was obtained from all participants for the publication of these case reports and any accompanying images. All research was carried out in accordance with relevant guidelines and regulations.

### iPSC culture and transfection

Human dermal fibroblasts were isolated from punch skin biopsies taken from three patients carrying biallelic mutations in the *SORD* gene (one patient with *SORD*^c.757delG/c.316_425+165del^, labelled SORD^-/-^#1; two sisters with *SORD*^c.757delG/c.757delG^, labelled SORD^-/-^ #2 and SORD^-/-^ #3), and two unaffected carriers with heterozygous mutations in SORD (*SORD*^c.757delG/wt^, labelled SORD^+/-^ #1; *SORD*^wt/c.316_425+165del^, labelled SORD^+/-^ #2), from two different families (**Figure 1A**). Fibroblasts obtained from a healthy, unrelated donor (labelled WT) were used as a control. Skin biopsies were conducted in accordance with the guidelines and with the approval of the Ethics Committee of the University of Naples “Federico II” (protocol number 334/21). Primary fibroblasts were obtained by cutting the biopsies into small pieces and culturing them in a fibroblast medium consisting of DMEM supplemented with 20% FBS, 2 mM L-glutamine, and 1% penicillin and streptomycin (all from Invitrogen), as previously described^20^. After 7–10 days of culture, the fibroblasts had spread and were routinely passaged using trypsin dissociation. For iPSC generation, we used a previously described protocol^21^. Briefly, 1 × 10⁶ fibroblasts were transfected with a total of 1 μg of the following plasmids, all from Addgene: pCXLE-hOCT3/4-shp53-F, pCXLE-hSK and pCXLE-hUL^22^, using a Nucleofector 2b device and an NHDF Nucleofector kit (both from Lonza), following the manufacturer’s instructions. After 21–24 days of reprogramming, individual iPSC clones were isolated by mechanical picking and expanded on Geltrex-coated plates in StemFlex medium (both from Thermo Fisher Scientific). They were then checked by immunofluorescence for positivity to the stemness markers OCT4 and NANOG. Three clones for each iPSC line were analysed for a normal karyotype. Karyotype analysis was performed by treating the iPSCs with 10 μg/ml colchicine overnight at 37 °C, followed by 20 minutes with 0.56% KCl. The iPSCs were then fixed in methanol and acetic acid (3:1), treated with trypsin (0.01% in PBS) and stained with 2% Giemsa at pH 6.8 (GTG banding). Analysis was performed using an ECLIPSE 1000 Nikon and GENIKON SYSTEM v3.9.8.

**Figure 1.**
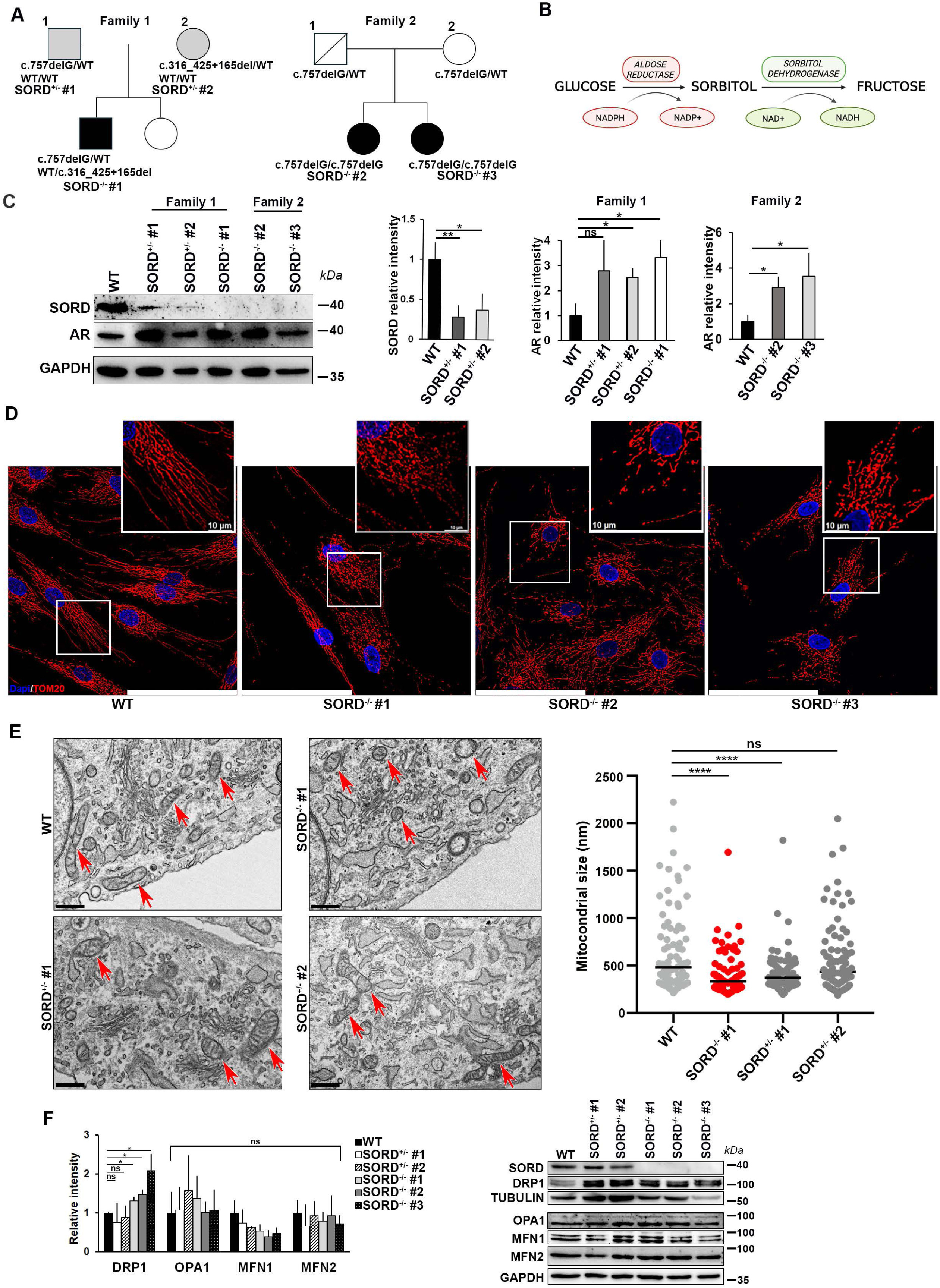
SORD deficiency alters mitochondrial morphology and remodelling in patient-derived fibroblasts. A) Representative pedigrees of individuals affected by CMT2/dHMN (black squares and circles) from two families that carry biallelic mutations in the *SORD* gene. Grey symbols indicate unaffected carriers in family 1 with heterozygous mutations in the *SORD* gene. In the text and in the Figures, the samples from individuals of the family 1 are referred to as follows: SORD^+/-^ #1 for carrier #1 with genotype *SORD* ^c.757delG/wt^, SORD^+/-^ #2 for carrier #2 with genotype *SORD* ^wt/c.316_425+165del^, SORD^-/-^#1 for patient #1 with genotype *SORD* ^c.757delG/c.316_425+165del^. In the text, the samples from individuals of the family 2 are referred to as follows: SORD^-/-^ #2 for patient #2 with genotype *SORD* ^c.757delG/c.757delG^, SORD^-/-^ #3 for patient #3 with genotype *SORD* ^c.757delG/c.757delG^. B) Overview of the polyol pathway, showing the two-step process involved in converting glucose to sorbitol (catalysed by the aldose reductase enzyme) and then to fructose (catalysed by the sorbitol dehydrogenase enzyme). C) Western blot analysis and relative quantification to evaluate SORD protein expression in lysates from unaffected carrier fibroblasts (SORD^+/-^, #1 and #2 from family 1) and in CMT2/SORD patient fibroblasts with biallelic *SORD* gene mutations (SORD^-/-^, #1 from family 1 and #2 and #3 from family 2) The data shown in the graphs are the mean ± SD (n = 3 biological replicates) of SORD and AR relative to total GAPDH signal intensity. Student’s t-test: ns, not significant; *p≤0.05, **p≤0.01). D) Fluorescence images of mitochondria stained with TOM20 (red) in WT and SORD-mutated fibroblast cells. The nuclei were stained with DAPI (blue), Scale bars = 100 µm. E) Transmission electron microscopy images of WT and SORD-mutated fibroblasts show that SORD-deficient fibroblasts have damaged, smaller mitochondria with an altered ultrastructure compared to WT cells (scale bars = 500 µm). Red arrows indicate mitochondria. The graph shows the quantification of the diameter of mitochondria in all the acquired images. Student’s t-test: ns, not significant; ****p≤0.0001. F) Western blotting images showing the expression profile of proteins involved in mitochondrial fission (DRP1) and fusion (OPA1, MFN1 and MFN2). The graph reports on the relative intensity as mean ± SD of the indicated proteins normalized against TUBULIN (for fission) or GAPDH (for fusion) as indicated. n=3 biological replicates. Student’s t-test: ns, not significant; *p≤0.05.

An unrelated, commercially available iPSC line (19-9-7T, WiCell Bank) was used as WT control. All iPSCs were routinely passaged in Stemflex medium on Geltrex-coated dishes.

### Mesodermal, ectodermal and endodermal differentiation of iPSCs

To assess their pluripotent potential, iPSCs obtained from different individuals were differentiated into derivatives of the three germ layers, as previously described^20,21,23^.

For mesodermal differentiation, undifferentiated iPSCs were seeded onto Geltrex-coated cell culture dishes at a density of 300,000 cells/cm² in StemFlex medium. After two days, differentiation was initiated by treating the cells with 12 µM CHIR99021 (Selleckchem) in RPMI (Gibco), supplemented with B27 minus insulin (Gibco), 2 mM L-glutamine, 0.1 mM 2-mercaptoethanol, non-essential amino acids, and penicillin–streptomycin (all from Invitrogen), for 24 h. The cells were then maintained in RPMI/B27-insulin medium for a further two days. On day 3, the cells were treated with 5 µM of the Wnt inhibitor IWP4 (Stemgent) in an RPMI/B27-insulin medium for 48 h. On day 5, the cells were washed once with RPMI to eliminate the inhibitor, after which they were maintained in an RPMI (Invitrogen) medium supplemented with B27 (Life Technologies), 2 mM L-glutamine, 0.1 mM 2-mercaptoethanol, non-essential amino acids, and penicillin-streptomycin (RPMI/B27 medium). From day 5 onwards, the cells were maintained in RPMI/B27 medium, with a medium change every two days. Beating cells appeared on days 14–16. For video recording, iPSC-derived cardiomyocyte monolayers were imaged at 37 °C in RPMI/B27 medium using LEICA DFC9000 GTC camera.

To induce endodermal differentiation, the iPSC clones were plated on Geltrex and cultured in RPMI medium supplemented with B27 and 100 ng/ml Activin A (Peprotech) for five days. The medium was then switched to RPMI supplemented with 20 ng/ml of BMP4 and 10 ng/ml of bFGF (both Preprotech) for five days. The cells were then grown for a further 5–7 days in Hepatocyte Culture Medium (Lonza), supplemented with 20 ng/ml of hepatocyte growth factor (Preprotech), followed by a final 5 days in RPMI-B27, supplemented with 20 ng/ml of hepatocyte growth factor (Preprotech).

See the next paragraph for ectodermal differentiation.

### Generation of MNPs and MNs from iPSCs and transfection

To generate a population of nearly pure MNPs, we adapted the protocol by *Du et al.* ^24^ as previously described^25^. Induced pluripotent stem cells (iPSCs) were dissociated with dispase and plated on Geltrex-coated plates at ∼20% confluence in Stemflex medium. After 24 hours, the medium was replaced with the following neural induction medium (NIM): DMEM/F12 and Neurobasal medium at a ratio of 1:1, supplemented with 1X N2, 1X B27 and 1X Glutamax (all from Invitrogen), 4 µM CHIR99021 (Selleckchem), 0.1 µM γ-secretase inhibitor XXI (Compound E, Millipore), 10 ng/ml human LIF (Millipore), and 3 µM SB431542 and 2 µM dorsomorphin (both Selleckchem). The medium was changed every two days for seven days. The differentiated cells were then dissociated using Accutase (Gibco) and split at a ratio of 1:3 in NIM supplemented with 10 ng/ml of human LIF (Millipore), 2 µM of SB431542 and 3 µM of CHIR99021. Once the cells were confluent, they were passaged onto Geltrex-coated plates in the same medium. After six passages, the cells were split 1:10 for a further 20 passages using Accutase in the same medium. At this point, the population of MNPs is positive for SOX1 and ISL1. These cells can be expanded and stored by freezing.

To obtain MNs, the MNPs were cultured in suspension in a 60 mm dish for one week in NIM supplemented with 0.1 mM ascorbic acid, 0.5 µM retinoic acid, and 0.1 µM purmorphamine to form 3D aggregates. The medium was changed every three days. The aggregates were then dissociated with trypsin into single cells and plated at a density of 0.5×10⁶ cells/ml in NIM supplemented with 10 µM Y-27632 (Sigma-Aldrich), 0.1 mM ascorbic acid (Sigma-Aldrich), 0.5 µM retinoic acid (Sigma-Aldrich), 0.1 µM purmorphamine (Sigma-Aldrich), 0.1 mM Compound E, 10 ng/ml GDNF and 10 ng/ml BDNF (both from Preprotech). The cells were cultured in these conditions for 10–12 days, with the medium changed every two days.

For siRNA transfection, 80% confluent MNPs were transfected with negative control or *SORD*-specific silencer siRNAs (both from Ambion,) using Lipofectamine Stem Reagent (Invitrogen), according to the manufacturer’s protocol. After 72 hours of transfection, the efficiency of the silencing was assessed by qPCR and Western blot.

### Sorbitol and ARi treatments

WT fibroblasts (2×10^5^ cells in p35 cell culture dishes) and neural precursors (3.5×10^5^ cells in geltrex-coated p35 cell culture dishes) were treated with different concentrations (0.1 and 0.5M) of sorbitol (Millipore) for 3 and 6 hours and maintained at 37 °C in a 5% CO2 humidified atmosphere.

For ARi-treatment, SORD-deficient fibroblasts and MNPs were treated with epalrestat (Merck) at concentration of 100 µM and 10 µM, based on previous reports^26,27^. For treatment of fibroblasts, ARi was added to the growth medium and left for 6 hours when the rescue of the normal mitochondrial network was already evident. To treat SORD-deficient MNs epalrestat was added to the growth medium on day 7 of differentiation and maintained for 3 days. In both cases, at the end of treatment, the cells were lysate or fixed to perform western blot or immunofluorescence.

### RNA extraction, reverse transcription and real time PCR

Total RNA was extracted from undifferentiated and differentiated iPSCs using Tri-Sure (Bioline), according to the manufacturer’s protocol One microgram of total RNA was used for reverse transcription with RevertAid Reverse Transcriptase (Thermo Scientific). Quantitative PCR (qPCR) was performed using a QuantStudio 7 Flex Real Time PCR System and Fast SYBR Green PCR Master Mix (Thermo Fisher Scientific). Gapdh mRNA expression was used as an internal control using the comparative 2^-ΔCt method. The gene-specific primers used are reported in Supplementary Table S1.

### Protein extraction and western blotting

For protein isolation, hDFs, iPSCs and MNPs were washed twice with ice-cold 1X phosphate saline buffer (1X PBS) and lysed with an extraction buffer containing 50 mM HEPES (Thermo Fisher Scientific), 150 mM NaCl (Applichem), 1 mM EDTA (pH8) (Applichem), 1 mM EGTA(pH8), 10% glycerol (Romil Pure Chemistry), and 1% Triton-X-100 (Applichem) complemented with protease (Sigma-Aldrich) and phosphatase inhibitors (PhosSTOP™, Roche), according to the manufacturer’s protocol. After lysis, the samples were centrifugated at 14000×g for 15 min at four degree to isolate total proteins. Concentration of total protein was determined using Bradford Protein assay (Bradford Reagent, Bio-Rad). Proteins were separated by SDS-PAGE under reducing conditions and blotted onto PVDF membrane (Merck Millipore). All the primary and HRP-conjugated secondary antibodies are listed in the **Supplementary Table 1**. Clarity Western ECL Substrate (Bio-Rad) was used for signals detection by enhanced chemiluminescence. Western blotting images were acquired with ChemiDoc MP Imaging System (Bio-Rad).

### Immunofluorescence analysis

Antibodies and materials used for immunofluorescence analysis are in Supplementary Table 1.

To assess alterations in the mitochondrial network, patient-derived fibroblasts, MNPs, and MNs were fixed in 4% paraformaldehyde for 10 minutes at room temperature and permeabilized with 0.05% saponin (Sigma-Aldrich) containing 1% bovine serum albumin for 15 minutes at room temperature, as previously described^25^. Cells were then incubated overnight at 4°C with primary antibodies. After washing with 1× PBS, cells were incubated for 1 hour at room temperature with secondary antibodies. Following additional washes, nuclei were stained with DAPI for 5 minutes at room temperature. Immunofluorescence images were acquired using a Leica Thunder Imaging System (Leica Microsystems) (Leica Microsystems) equipped with a LEICA DFC9000 GTC camera, lumencor fluorescence LED light source, unless otherwise specified. To analyse the mitochondrial network in MNPs upon *SORD* silencing, images were acquired using a Zeiss LSM980 confocal microscope.

For morphometric analysis skin-derived fibroblasts were plated on 8-well chambers (Ibidi) and stained with 50 nM MitoTracker™ Green FM (Thermo Fisher Scientific) for 30 minutes at 37°C in a humidified 5% CO₂ atmosphere to quantify mitochondrial defects. After staining, cells were washed with 1× PBS and maintained in growth medium. Live z-stack images and time-lapse recordings were acquired using a Zeiss LSM980 confocal microscope.

To assess stemness and differentiation marker expression, undifferentiated or differentiated iPSC, MNPs and MNs were fixed with 4% paraformaldehyde, permeabilized with 0.5% Triton X-100, and blocked in 10% FBS and 1% BSA in 1× PBS, as previously described^21^. Clones were incubated overnight at 4°C with primary antibodies, followed by incubation with secondary antibodies (all listed in **Supplementary Table 1**). Nuclei were stained with DAPI for 5 minutes at room temperature. The number of SOX1/ISL1-positive MNPs and ChAT and ISL1 MNs was determined by acquiring ten different images for each sample (WT, SORD^+/-^#1 and #2, SORD^-/-^#1) with an average of about 130 cells identified with DAPI per image. Counting was carried out in biological triplicate.

To evaluate sorbitol-induced cell death, treated fibroblasts and MNPs were fixed with 4% paraformaldehyde and permeabilized with saponin, as described above. Cells were incubated overnight at 4°C with anti-cleaved caspase-3 (Asp175) and anti-TOM20 primary antibody, followed by incubation with secondary antibodies as described above. Nuclei were stained with DAPI for 5 minutes at room temperature. To determine the number of cleaved caspase-3–positive cells in sorbitol-treated hDFs and MNPs, images of at least 350 cells identified by DAPI staining were acquired from three independent experiments.

To quantify reactive oxygen species levels, sorbitol-treated fibroblasts and MNPs were incubated with dihydroethidium (DHE) (Thermo Fisher Scientific) at a final concentration of 25 µM for 1 hour at 37°C in a humidified 5% CO₂ atmosphere. After staining, cells were washed with 1× PBS and fixed with 4% paraformaldehyde for 10 minutes at room temperature. DHE fluorescence signal was quantified as integrated density/nuclear area using Fiji software as integrated density normalized to nuclear area. The quantification of DHE signal was carried out by analyzing a total number of about 100 cells for hDFs 200 cells for MNPs from biological replicates (n=2). Mean DHE intensity was calculated considering all analyzed cells, and data are expressed as mean ± SEM. Statistical comparisons between groups were performed using an unpaired Student’s t-test.

### Mitochondrial network analysis

Image processing and quantitative analysis were performed using ImageJ with a custom macro^25^. Multichannel microscopy datasets were imported via the Bio-Formats Importer, and Z-stacks were reduced to two-dimensional representations through maximum intensity projection. Images were subsequently converted to 8-bit grayscale and denoised using a Gaussian blur filter (σ = 1). Mitochondrial structures were segmented using the appropriate automatic thresholding algorithm under a dark background assumption and individual objects were detected using the Analyze Particles function. Morphometric descriptors, including area, perimeter, circularity and Feret’s diameter, were extracted. In parallel, binary masks were skeletonized and processed with the Analyze Skeleton (2D/3D) plugin to quantify network topology parameters, such as branch length. The analysis pipeline generated both annotated skeleton images and tabulated summary outputs^29^.

Data visualization and statistical analyses were performed with R software, using the packages grid, ggplot2 and ggpubr, as reported in the Supplementary Table 1.

### Neurite network analysis

Images were analyzed using a two-step automated workflow combining ImageJ/Fiji preprocessing and morphometric quantification with R statistical analysis^28^. Multichannel microscopy images were batch-processed by channel separation, background subtraction, Gaussian filtering, intensity normalization and 8-bit conversion. Cytoplasmic signal was then projected (maximum intensity projection), thresholded to generate binary masks, skeletonized and analyzed using the Analyze Skeleton plugin to extract neurite structural parameters^29^. In addition, distance-map based measurements were used to estimate branch and junction spatial density. Data visualization and statistical analyses were performed using the packages grid, ggplot2, and ggpubr, as reported in the Supplementary Table 1.

### Cell metabolism analysis

Real-time measurements of oxygen consumption rate (OCR) were made using an XF Extracellular Flux Analyzer (Agilent Technologies, Santa Clara, CA). The day before the experiment cells were plated on 96-well SeaHorse cell culture plates (103794 Agilent) in culture medium (100,000 cells/well). Cells were incubated overnight before cell metabolism assay.

The day of the experiment, cells were washed once and kept in bicarbonate-free XF DMEM medium pH7.4 (Agilent) supplemented with 1 mM pyruvate, 2 mM glutamine and 5 mM glucose. The cells were incubated for 1h in a 37°C non-CO2 incubator. Mitochondrial respiration was monitored at basal state and after sequential injection of the mitochondrial modulators oligomycin (3 µM), FCCP (6 µM) and antimycin/Rotenone (2.5 µM and 1 µM respectively) (all chemicals from Sigma Aldrich) that induce mitochondrial stress^30^. After the assay, cells were lysed in RIPA buffer, total protein content was quantified by BCA assay (Pierce) and the value was used for normalization of the OCR values.

### Electron microscopy studies

HDFs or MNs were fixed with 1% glutaraldehyde prepared in 0.2 M HEPES buffer (pH 7.3) for 30 min at room temperature. Samples were then post-fixed in OsO4 and uranyl acetate, dehydrated, embedded in EPON resin and polymerized at 60°C for 72 h, as previously described^31^. Ultrathin sections (60 nm) were cut using a Leica EM UC7 microtome. Electron microscopy images were acquired using a FEI Tecnai-12 transmission electron microscope (Thermo Fisher) equipped with a VELETTA CCD digital camera. The mitochondrial diameter was measured using Fiji software.

### Statistical analysis

Statistical analyses reported in this study were performed using Microsoft Excel or GraphPad Prism (version 7.0a). Quantitative differences between experimental groups were evaluated using unpaired, two-tailed Student’s t-tests, unless otherwise specified. Statistical significance was set at p value at least ≤0.05. Data are presented as mean ± standard deviation, unless otherwise specified.

Biological replicates were defined as independent experimental repeats. All the output datasets for the morphometric analysis of the mitochondrial and neurite networks were compiled across experimental groups and statistically compared with the control samples using unpaired *t*-tests. Results were visualized as boxplots, where the horizontal line within each box indicates the median value, while the upper and lower borders show the interquartile range (IQR). Whiskers extend to the 5th and 95th percentiles, providing a representation of data dispersion beyond the central distribution.

## Results

### *SORD* mutations alter the expression of polyol pathway enzymes

To investigate the functional effects of SORD deficiency, we recruited three patients carrying different combinations of *SORD* mutant alleles from two families, along with two unaffected carriers (the parents of patient #1, family 1), all previously described in *Cortese et al.*^7^ (**Figure 1A**). The affected individuals exhibited distinct clinical phenotypes. Patient #1 (hereafter referred as SORD^-/-^#1), who carried compound heterozygous mutations (*SORD*^c.757delG/c.316_425+165del^; family 1, **Figure 1A**), presented with axonal sensorimotor neuropathy, with disease onset during adolescence^7^. His parents were unaffected heterozygous carriers (*SORD*^c.757delG/wt^, hereafter indicated as SORD^+/-^ #1, and *SORD*^wt/c.316_425+165del^, hereafter indicated as SORD^+/-^ #2) (**Figure 1A**). The two affected sisters from family 2 (hereafter indicated as SORD^-/-^ #2 and SORD^-/-^ #3) carried the common homozygous mutation (*SORD*^c.757delG/c.757delG^) and developed a late-onset disease (around their 40s) characterized by walking difficulties. Nerve conduction studies revealed distal hereditary motor neuropathy (dHMN) without sensory deficits^7^. None of the three patients showed evidence of other causes of acquired neuropathy, including diabetes.

Primary dermal fibroblasts were isolated from punch skin biopsies of the affected individuals with CMT2/SORD, their familial controls (described above), and an unrelated healthy subject (hereafter indicated as WT). We then analyzed the levels of the two enzymes involved in the polyol pathway, (**Figure 1B**). We first assessed SORD protein levels and found that both biallelic mutations resulted in complete loss of SORD protein in the fibroblasts from all three patients (SORD^-/-^ #1, SORD^-/-^ #2 and SORD^-/-^ #3). In contrast, heterozygous carriers (SORD^+/-^ #1 and SORD^+/-^ #2) exhibited an approximately 50% reduction in SORD protein levels (**Figure 1C**), consistent with previous findings^7^. Then, we investigated whether SORD deficiency affects the levels of AR protein, the rate-limiting enzyme of the polyol pathway^32^. Western blot analysis of protein extracts from both WT and SORD-mutant fibroblasts revealed a significant accumulation of AR protein in all three SORD^−/−^ affected individuals compared to the WT control (**Figure 1C**). The findings, taken together, indicate that SORD mutations result in the loss of SORD protein, which is associated with dysregulation of AR expression. This, in turn, suggests alterations at both steps of the polyol pathway.

### SORD deficiency affects mitochondrial morphology and network architecture

Mitochondrial dysfunction is a common feature across multiple CMT2 subtypes^17,18,33^, and data from *Drosophila* indicated that also *SORD* mutation can lead to mitochondrial dysfunction^15^. Thus, we investigated whether patient cells exhibited alterations in the mitochondrial compartment. To this end, we performed immunofluorescence analyses on control (WT), carrier (SORD^+/-^ #1 and SORD^+/-^#2), and patient (SORD^-/-^ #1, SORD^-/-^ #2 and SORD^-/-^ #3) -derived fibroblasts using an antibody against the outer mitochondrial membrane protein TOM20. Our analysis revealed that SORD-deficient cells lost the typical filamentous and interconnected mitochondrial network characteristic of fibroblasts (**Figure 1D**). Indeed, patient cells displayed smaller mitochondria and a less complex network compared to control cells (**Figure 1D**). No significant changes in mitochondrial morphology were observed in fibroblasts from unaffected heterozygous carriers (**Figure S1A**).

Mitochondrial structural alterations were further characterized using MitoTracker™ Green FM, a fluorescent dye that labels mitochondria independently of membrane potential^34^ (**Supplementary Movies**). Quantitative analysis revealed a marked reduction in mitochondrial area and perimeter in SORD-deficient cells (**Figure S1B**), along with increased fragmentation compared to control cells. While the number of branches remained unchanged, patient cells exhibited significantly shorter average branch lengths and reduced individual mitochondrial branch length (**Figure S1B**), indicative of a smaller, more fragmented, and less connected mitochondrial network.

These observations were further supported by transmission electron microscopy (TEM) analysis. Morphometric assessment of TEM images confirmed that mitochondria in patient cells were smaller and exhibited disrupted cristae structure, in contrast to the well-preserved ultrastructure observed in control cells (**Figure 1E**). Notably, mitochondrial morphology and cristae architecture in carrier fibroblasts were comparable to those of unrelated control (**Figure 1E**).

Alterations in mitochondrial network architecture are often associated with dysregulation of mitochondrial dynamics. Therefore, we next examined the expression of key proteins involved in mitochondrial fission and fusion. Notably, loss of SORD resulted in increased levels of dynamin-related protein 1 (DRP1), a GTPase that mediates mitochondrial fission (**Figure 1F**). In contrast, no significant changes were observed in the expression of fusion-related proteins, including mitofusin 1 and 2 (MFN1 and MFN2), or optic atrophy 1 (OPA1) (**Figure 1F**).

Collectively, these findings indicate that SORD deficiency disrupts mitochondrial network organization, that can be due from an imbalance in mitochondrial dynamics, likely reflecting a shift toward increased fission activity.

### SORD deficiency alters mitochondrial respiration and induces mitochondrial damage

MNs are the primary cell type affected in CMT2-SORD disease. However, to date, a well-characterized human model that faithfully recapitulates the disease phenotype in MNs has been lacking. To address this, we generated an *in vitro* model of CMT2/SORD MNs by reprogramming carrier and patient fibroblasts (*SORD*^c.757delG/wt^, *SORD*^wt/c.316_425+165del^, and *SORD*^c.757delG/c.316_425+165del^, indicated as SORD^+/-^ #1, SORD^+/-^ #2 and SORD^-/-^ #1, respectively) into induced pluripotent stem cells (iPSCs). An unrelated iPSC cell line was used as WT control.

Selected iPSC clones were validated for normal karyotype and sustained expression of pluripotency markers NANOG and OCT4 after extended culture (≥15 passages) (**Figure S2A,B**). Western blot analysis confirmed the absence of SORD protein in patient-derived iPSCs (**Figure S2C**). Pluripotency was further verified by differentiation into derivatives of the three germ layers (**Figure S2D**).

To generate MNs from control, carrier, and patient iPSCs, we used a slightly modified version of a previously established protocol^24^. This approach includes an initial step yielding highly enriched (>95%) motor neuron progenitors (MNPs) that can be expanded, followed by a second differentiation step to obtain a population enriched in MNs (>90%). Proper differentiation into MNPs was confirmed by expression of the neural progenitor marker SOX1 and the MN lineage marker ISL1 (**Figure S3A**). To assess the impact of SORD deficiency in neural cells, we first examined mitochondrial morphology in control and patient-derived MNPs. By immunofluorescence analysis we found that SORD-deficient MNPs displayed clear mitochondrial fragmentation, evidenced by the presence of rounded mitochondria (**Figure 2A, white arrows**). This phenotype was associated with enhanced mitochondrial fission, as indicated by elevated levels of DRP1 and phosphorylated DRP1^S616 in patient MNPs relative to WT cells (**Figure 2B**). In contrast, no changes were observed in the expression of the fusion marker MFN2, in agreement with the phenotype observed in fibroblasts (**Figure 1F**). An increase in DRP1 levels was also detected in heterozygous MNPs carrying the *SORD*^wt/c.316_425+165del^ ^mutation^ (SORD^+/-^ #2), although phospho-DRP1^S616 levels were unchanged. This suggests that the *SORD*^wt/c.316_425+165del^ mutation (SORD^+/-^ #2) may exert a stronger effect than the *SORD*^c.757delG/wt^ mutation (SORD^+/-^ #1); however, increased DRP1 alone was insufficient to induce evident mitochondrial structural alterations in heterozygous cells (**Figure 2A**). Consistent with observations in fibroblasts (**Figure 1C**), SORD-deficient MNPs also exhibited accumulation of AR protein (**Figure 2B**), indicating that this response is a general consequence of SORD deficiency. To confirm that the mitochondrial phenotype is specifically linked to SORD loss and not limited to patient-derived cells, we performed SORD knockdown in WT MNPs using a specific siRNA against *SORD* mRNA (**Figure S3B**). The reduction of SORD protein (∼40%), accompanied by AR accumulation (**Figure S3C**), recapitulated the mitochondrial defects observed in SORD−/− MNPs (**Figure 2C**). Quantitative analysis revealed increased mitochondrial circularity and reduced aspect ratio and branch length (**Figure 2D** and **S3D**), consistent with enhanced fragmentation.

**Figure 2.**
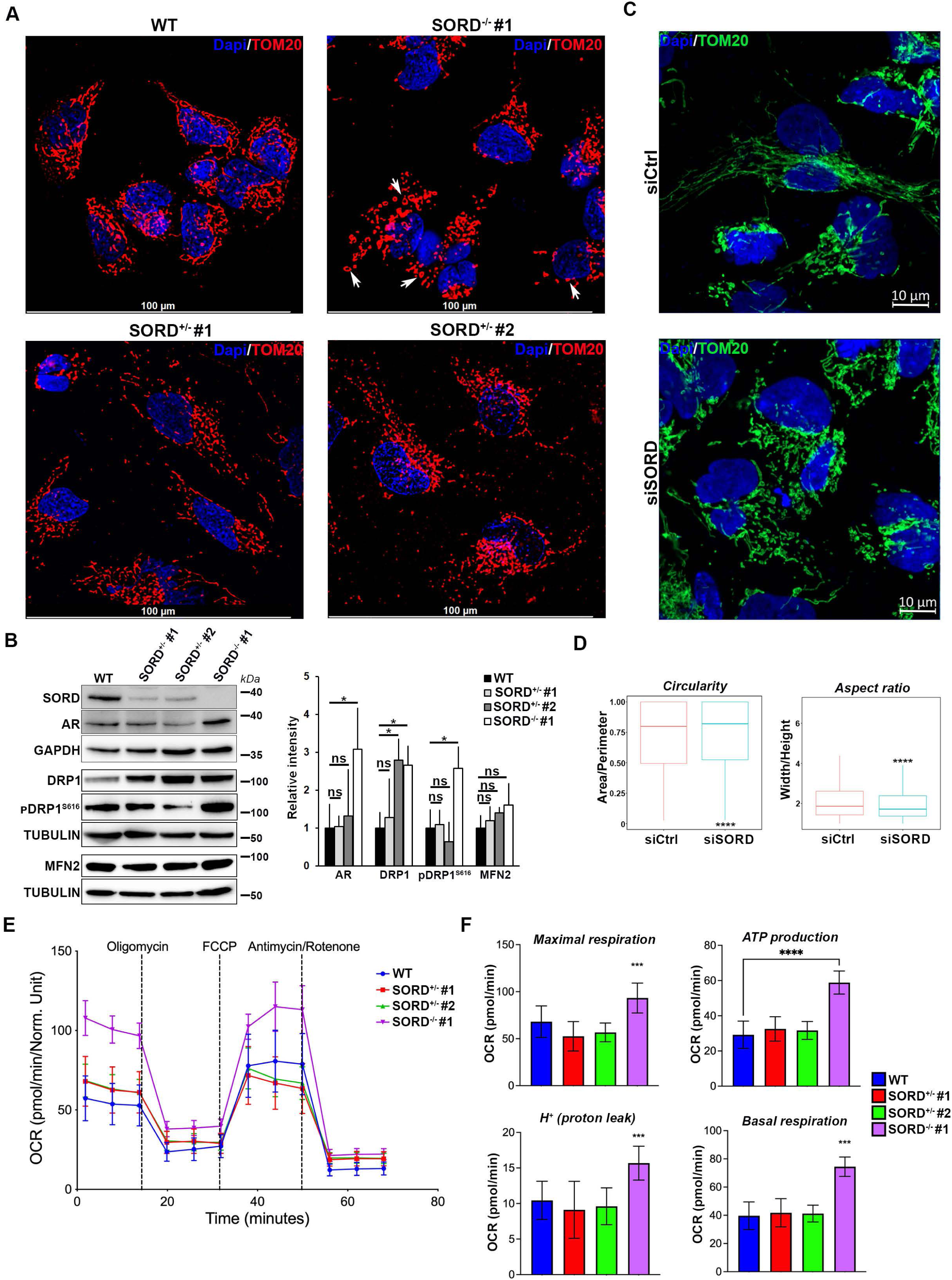
SORD deficiency increases mitochondrial fission in neural cells. A) Immunofluorescence analysis of MNPs differentiated from WT, carriers (SORD^+/-^ #1 and SORD^-/+^ #2) and patient-derived (SORD^-/-^ #1) iPSCs. Mitochondrial network was stained using TOM20 antibody (red). White arrows indicate rounded mitochondria present in patient cells but absent in control and carrier cells. Scale bars = 100 µm. B) Western blotting images and the relative quantification of the mitochondrial fission proteins DRP1 and pDRP1S616, as well as the fusion protein MFN2 in SORD-deficient MNPs compared to controls. The graph shows the mean ± SD relative intensity of the indicated proteins, normalized to total GAPDH (for AR) and TUBULIN (for all others), for three biological replicates (n=3). Student’s t-test: ns, not significant; *p≤0.05. C) Confocal images to analyze mitochondrial network in MNPs transfected with a control siRNA (siCtrl) and a specific *SORD* siRNA (si*SORD*). Scale bars = 10 µm. D) Morphometric analysis of mitochondria in MNPs transfected with control and *SORD* specific siRNAs. The graphs represent mitochondrial circularity (area/perimeter) and the aspect ratio (width/height) in the indicated samples. Box plots indicate the interquartile range (IQR, 25th–75th percentile), with the horizontal line indicating the median. The whiskers extend to the minimum and maximum values, and red triangles denote the mean for each group. Student’s t-test: ns, not significant; ****p≤0.0001). E) Cell metabolism analysis showing the oxygen consumption rate (OCR) in basal conditions and after the sequential injection of the mitochondrial modulators in WT and SORD-deficient MNPs. F) The graphs show the evaluation of maximal respiration, ATP production, proton leak and basal respiration as mean ± SD of three biological replicates (n=3). Student’s t-test: ns, not significant; ***p≤0.0005, ****p≤0.00005).

Given that alterations in mitochondrial network are often associated with metabolic dysfunction, we next analyzed mitochondrial metabolism. MNPs were used for these experiments due to their suitability for metabolic measurements. Oxygen consumption rate (OCR) and extracellular acidification rate (ECAR) were measured in live control and patient MNPs (see Methods). During the assay, oligomycin, FCCP (carbonyl cyanide-4-(trifluoromethoxy) phenylhydrazone), and antimycin A/rotenone were sequentially added under basal conditions. Oligomycin inhibits ATP synthase, allowing measurement of ATP-linked respiration and proton leak. FCCP uncouples oxidative phosphorylation by dissipating the mitochondrial membrane potential, enabling assessment of maximal respiratory capacity. Antimycin A and rotenone inhibit the electron transport chain, allowing quantification of non-mitochondrial respiration. Patient-derived MNPs exhibited a marked increase in OCR compared to controls (**Figure 2E,F**), indicating enhanced oxidative phosphorylation and suggesting higher basal energetic demand. These observations, together with mitochondrial fragmentation, are commonly associated with metabolic stress^35,36^. Maximal respiration (after FCCP treatment) was also increased, indicating that SORD-deficient cells retain the ability to respond to metabolic stress. Despite increased ATP production, patient MNPs showed a tendency toward elevated proton leak, reflecting a component of basal respiration that is uncoupled from ATP synthesis (**Figure 2F**). This feature is widely considered a hallmark of mitochondrial dysfunction^37^.

### Mitochondrial and neurite networks are altered in SORD-deficient neural cells

To investigate how the alterations observed in patient fibroblasts and MNPs relate to peripheral neuron degeneration in CMT2/SORD, we differentiated control and patient-derived MNPs into MNs. Given that neurite network defects have been reported in iPSC-derived neuronal models of various CMT2^33^, we performed immunofluorescence analyses using an antibody against the neuronal cytoskeletal marker β3-TUBULIN. Control and carrier MNs formed a highly interconnected network, characterized by dense neurite branching and numerous junctions (**Figure 3A**). In contrast, patient-derived MNs exhibited marked structural abnormalities in neuronal network organization (**Figure 3A**). These defects were associated with impaired neuronal morphology and reduced connectivity, as SORD-deficient MNs formed fewer intercellular connections and displayed defective neurite outgrowth (**Figure 3B**). Importantly, these abnormalities were not due to impaired differentiation, as the proportion of MNs expressing intermediate and mature MN markers (ISL1 and CHAT) was comparable between patient and control cells (**Figure 3A** and **S4A**). To quantitatively assess these structural defects, we performed dimensional analysis of the neuronal network. SORD-deficient MNs exhibited impaired branching, with a significant reduction in neurite branch length compared to WT (**Figure 3B**). In addition, patient MNs showed increased branch distance despite having a similar number of branches and junctions (**Figure 3B**), suggesting a sparse and less efficiently connected neuronal network. These results indicated that SORD deficiency negatively affects the neurite network of iPSC-derived MNs.

**Figure 3.**
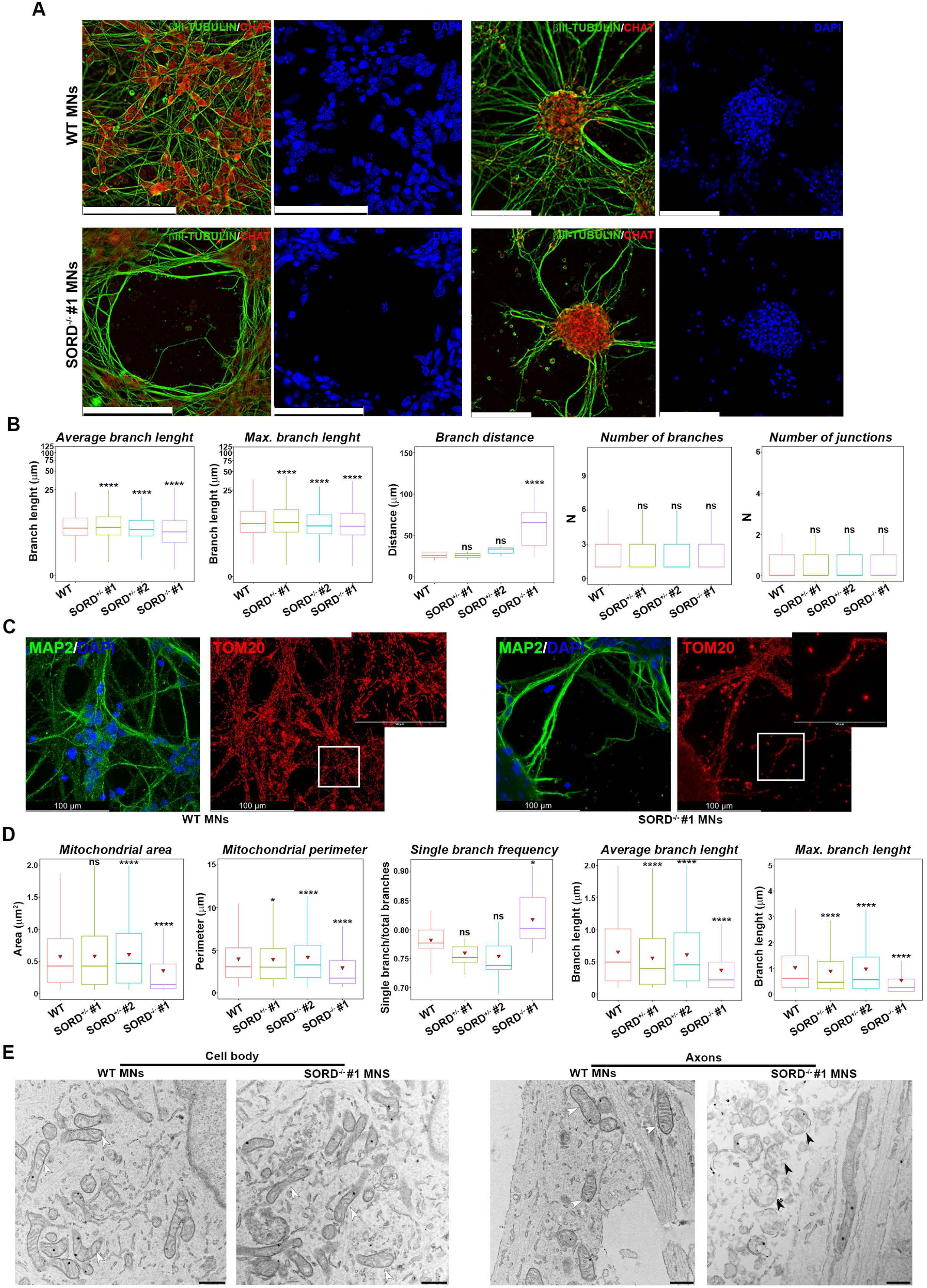
SORD-deficient MNs exhibit altered neurite and mitochondrial networks. A) Immunofluorescence analysis of WT and SORD-deficient MNs. The neuronal cytoskeleton was stained with βIII-TUBULIN (green) and CHAT (red) was used as a marker of MNs. Two representative images are shown for each condition. Scale bar = 100 µm. B) Morphometric analysis of neuronal cytoskeleton of WT, carriers and patient MNs to evaluate average and maximum branch length (µm) of neurites and the distance (µm) between neurites. Box plots indicate the interquartile range (IQR, 25th–75th percentile), with the horizontal line indicating the median. The whiskers extend to the minimum and maximum values, and red triangles denote the mean for each group. Student’s t-test: ns, not significant; ****p ≤ 0.0001). C) Immunofluorescence analysis to evaluate the mitochondrial network of WT and SORD-deficient MNs. TOM20 (red) was used to stain the mitochondria, MAP2 (green) to stain the neuronal microtubules, and DAPI (blue) to stain the nuclei. Scale bars = 100 µm. The white square indicates the cropped region. D) Morphometric analysis of mitochondria in MNs from WT, carriers (SORD^+/-^ #1, SORD^+/-^ #2) and patient (SORD^-/-^ #1). The graphs report on dimensions (area and perimeter) of the mitochondria and structure of the altered mitochondrial network (single and average branch length). Box plots indicate the interquartile range (IQR, 25th–75th percentile), with the horizontal line indicating the median. The whiskers extend to the minimum and maximum values, and red triangles denote the mean for each group. Student’s t-test: ns, not significant; *p≤0.05, ****p ≤ 0.0001). E) Transmission electron microscopy to evaluate the structure of mitochondria in the cell bodies and axons of WT and patient MNs. The white arrowheads indicate normal mitochondria, while the black arrowheads indicate damaged axonal mitochondria in SORD-deficient MNs. Scale bars: cell body=600nm, axons=500nm.

We next analyzed the mitochondrial network in MNs using immunofluorescence and quantitative image analysis. Consistent with findings in fibroblasts and MNPs, patient-derived MNs exhibited pronounced mitochondrial alterations (**Figure 3C**). Specifically, mitochondrial area and perimeter were reduced, and the mitochondrial network lost its typical interconnected organization, with an increased frequency of isolated mitochondrial branches (**Figure 3D**). Furthermore, mitochondrial branches were significantly shorter, as indicated by reduced average and maximum branch lengths compared to WT and heterozygous MNs (**Figure 3D**).

Given that SORD-deficient MNs exhibit both neurite and mitochondrial abnormalities, we performed transmission electron microscopy (TEM) to determine whether mitochondrial defects were present in both the cell bodies and neurites. TEM analysis revealed severe structural damage in axonal mitochondria, whereas mitochondria within the cell body appeared largely preserved and comparable to those in WT cells (**Figure 3E**). These observations can explain why SORD mutations exert particularly severe effects on neural cells. Thus, to understand whether these defects are accompanied by alteration in cell survival, we evaluated apoptosis in MNs by analyzing caspase-3 activation. Immunofluorescence analysis showed a significantly higher number of cells positive for active caspase-3 in patient-derived MNs compared to WT control (**Figure S4B**).

Taken together, these findings demonstrate that SORD-deficient MNs exhibit severe defects in both neurite and mitochondrial network organization and are more prone to cell death, consistent with the neurodegenerative phenotype observed in patients.

### Sorbitol accumulation increases AR expression and induces oxidative stress

*Cortese and colleagues* previously demonstrated that fibroblasts derived from patients with biallelic *SORD* mutations accumulate intracellular sorbitol^7^. Because sorbitol accumulation can disrupt membrane permeability, alter osmotic balance, and induce oxidative stress^38^, we ask whether the mitochondrial fragmentation observed in SORD-deficient cells can be mimicked by treatment with sorbitol. To this end, WT fibroblasts were treated for 6 hours with sorbitol at concentrations of 0.1 M and 0.5 M, exceeding the physiological intracellular range (0.01–0.05 M) reported previously^39^. We found that sorbitol treatment induced mitochondrial fragmentation, with more pronounced effects at higher concentrations, closely resembling the phenotype observed in SORD-deficient cells (**Figure 4A**). Quantitative analysis of confocal images revealed significant reductions in mitochondrial area and perimeter (**Figure 4B**), along with marked disruption of the mitochondrial network. In particular, mitochondrial branch length was significantly decreased in sorbitol-treated fibroblasts compared to untreated cells (**Figure 4B** and **S5A**). Additionally, a reduction in mitochondrial aspect ratio (length/width) and an increase in mitochondrial circularity were observed (**Figure 4B**), consistent with enhanced mitochondrial fission and fragmentation^40^.

**Figure 4.**
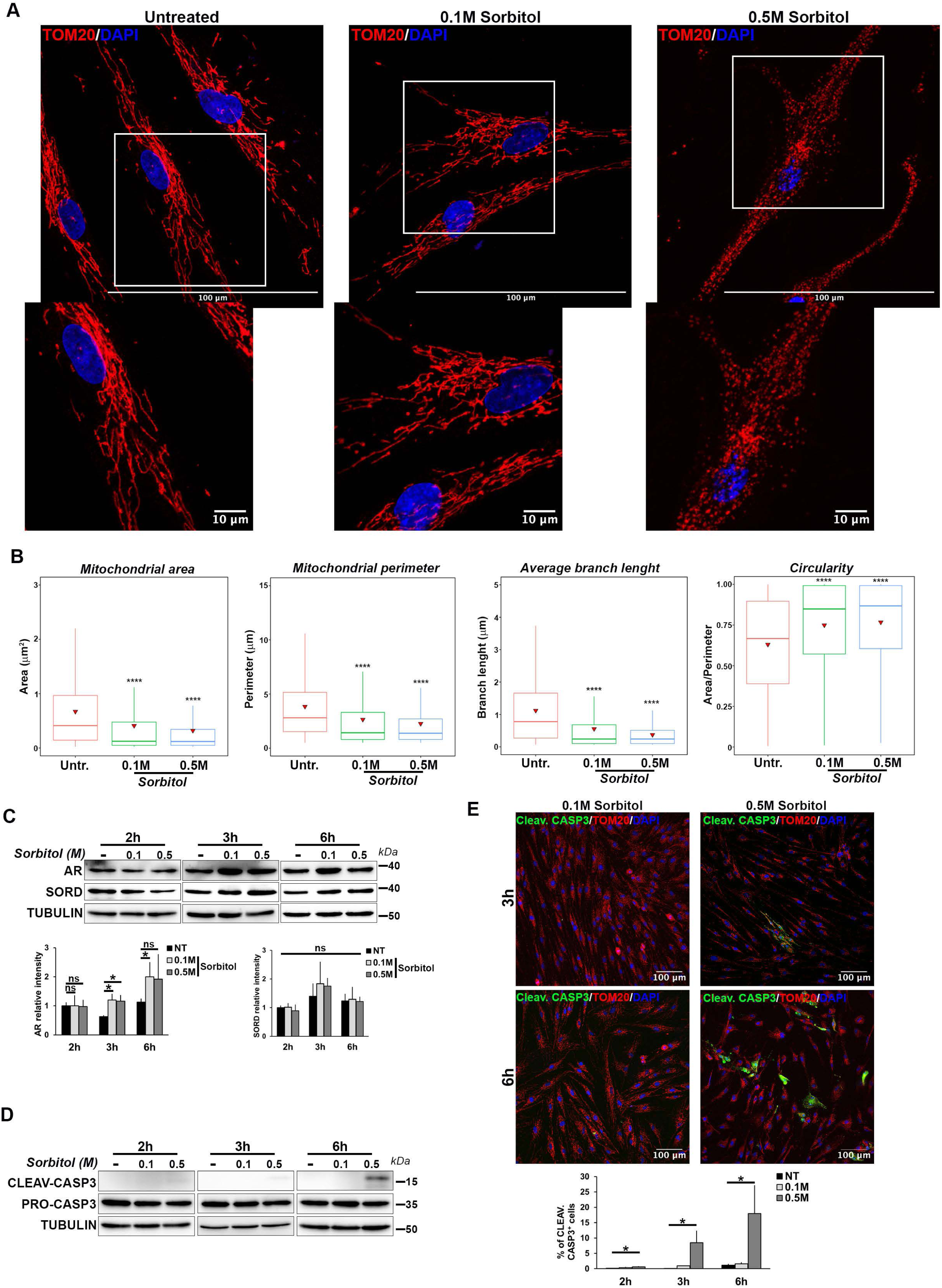
Sorbitol treatment induces mitochondrial fragmentation and cell death. A) Immunofluorescence analysis of mitochondria in fibroblasts untreated or treated with two concentrations of sorbitol (0.1 and 0.5 M). TOM20 (red) was used to stain the mitochondria and DAPI (blue) to stain the nuclei. Scale bars = 100 µm. Cropped images were shown for each condition. B) Morphometric analysis of mitochondria in fibroblasts untreated or treated with the indicated concentrations of sorbitol. The graphs report the analysis of mitochondrial dimensions (area and perimeter) and mitochondrial network parameters (average branch length and circularity). Box plots indicate the interquartile range (IQR, 25th–75th percentile), with the horizontal line indicating the median. The whiskers extend to the minimum and maximum values, and red triangles denote the mean for each group. Student’s t-test: ns, not significant; ****p≤0.0001). C) Western blot analysis and relative quantification of AR protein expression after sorbitol treatment. The data in the graphs represent the mean ± SD of SORD and AR relative to total TUBULIN signal intensity of three biological replicates (n=3). Student’s t-test: ns, not significant; *p≤0.05). D) Western blot analysis of CLEAVED and PRO-CASPASE 3 at different time points of sorbitol treatment. PRO-CASPASE 3 and TUBULIN were used as internal controls. E) Immunofluorescence analysis to detect fibroblasts positive for active CASPASE 3 upon treatment with different concentrations of sorbitol (0.1 and 0.5 M) for three and six hours. CLEAVED CASPASE 3 (green) was used as marker of apoptosis; TOM20 (red) was used to stain mitochondria. Scale bars = 100 µm. The graph shows the mean ± SD (n = 3 biological replicates) of the percentage of fibroblasts positive for CLEAVED CASPASE 3 in the conditions indicated. Student’s t-test: ns, not significant; *p≤0.05.

We next investigated whether sorbitol accumulation can lead to accumulation of AR protein. WT fibroblasts treated with sorbitol at concentrations of 0.1 M and 0.5 M exhibited elevated AR protein levels under both conditions (**Figure 4C**). However, statistically significant increase in AR expression following prolonged treatment (6 hours) was observed only at the lower sorbitol concentration (0.1 M). This plateau is likely due to cytotoxic effects at high sorbitol doses, which activate cell death pathways, as indicated by increased levels of cleaved caspase-3 (**Figure 4D**). Immunofluorescence analysis and relative quantitation confirmed activation of cleaved caspase-3 under these conditions (**Figure 4E**), in agreement with previous reports showing that sorbitol induces apoptosis via caspase-3 activation^15^.

Since AR is accumulated upon sorbitol treatment and increased AR activity has been associated with oxidative stress^41^, we evaluated reactive oxygen species (ROS) production after sorbitol treatment. Staining with dihydroethidium (DHE), a fluorescent ROS detector, an increase in ROS levels was quantified in sorbitol-treated cells compared to the untreated condition (**Figure S5B**). These results indicate that sorbitol treatment leads to both AR accumulation and oxidative stress.

Sorbitol accumulation via the polyol pathway has been proposed as a key mechanism underlying peripheral neuropathies associated with altered glucose metabolism^14,42^, suggesting that neural cells may be particularly vulnerable. To test this hypothesis, WT MNPs were treated with sorbitol under the same conditions used for fibroblasts. Notably, MNPs were more sensitive to sorbitol-induced stress than fibroblasts. After prolonged exposure (6 hours) to high sorbitol concentrations (0.5 M), MNP morphology was severely affected, with detectable alterations already at 0.1 M, whereas fibroblasts under the same conditions remained viable and showed only mild stress (**Figure S5C**). This increased sensitivity was further supported by the observation that alterations in mitochondrial network of MNPs treated with 0.1M of sorbitol were observed as early as 3 hours of treatment (**Figure 5A**). Quantitative analysis highlighted reductions in mitochondrial area, perimeter, and branch length, accompanied by increased circularity, particularly in MNPs treated with 0.5 M sorbitol (**Figure 5B**). In parallel, sorbitol treatment induced an early and dose-dependent increase in ROS levels in MNPs (**Figure S5D**).

**Figure 5.**
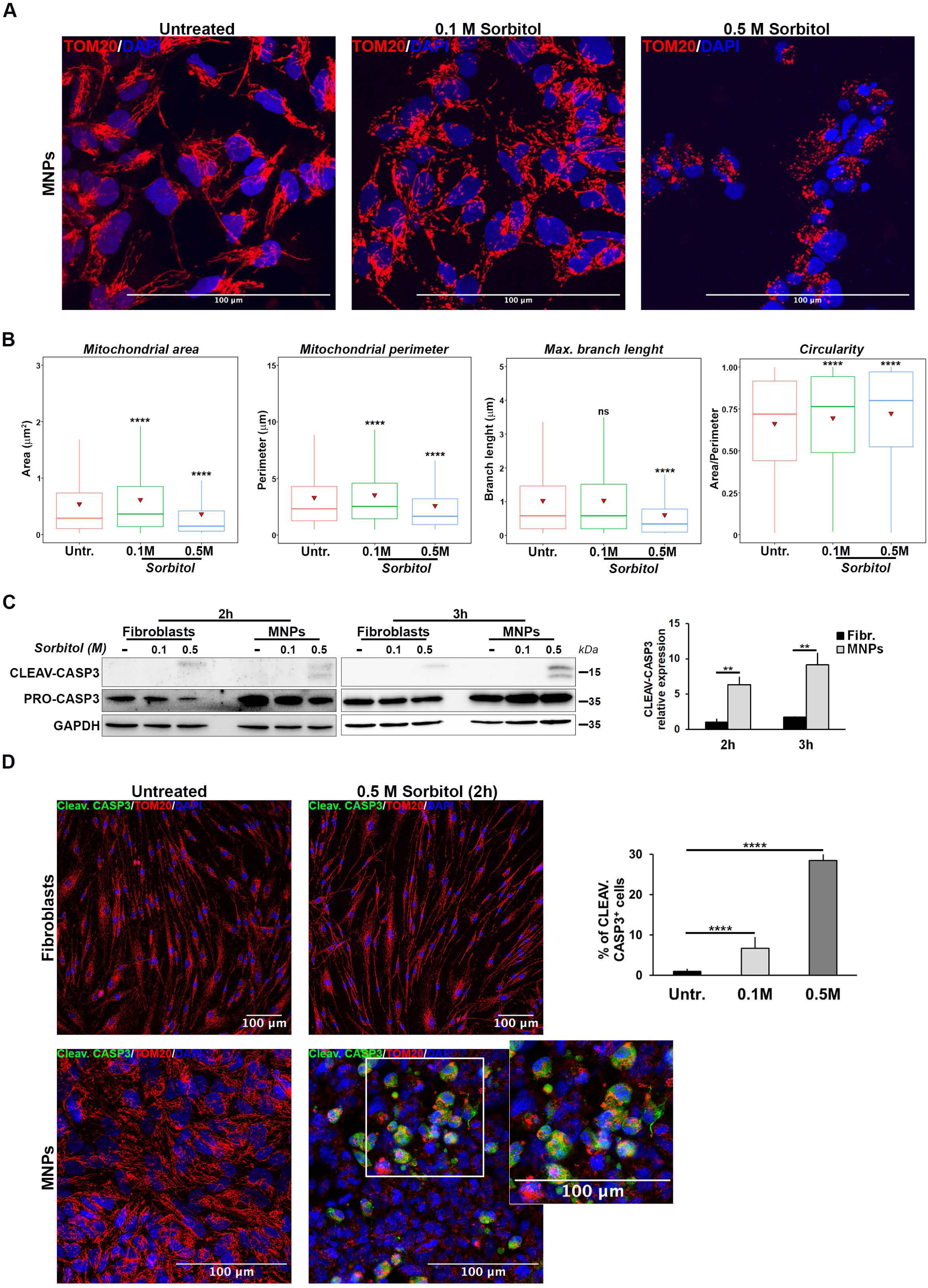
MNPs are more sensitive than fibroblasts to sorbitol overload. A) Confocal images showing mitochondrial network in MNPs treated with the indicated concentrations of sorbitol for three hours. Scale bars = 10 µm. B) Morphometric analysis of the immunofluorescence in the panel (A). Box plots reports on the analysis of mitochondrial area, perimeter, branch length and mitochondrial circularity. The values are presented as the interquartile range (IQR, 25th–75th percentile), with the horizontal line indicating the median. Whiskers extend to the minimum and maximum values, and red triangles denote the mean for each group. Student’s t-test: ns, not significant; ****p≤0.0001). C) Western blot analysis of fibroblasts and MNPs treated with the indicated concentrations of sorbitol for two and three hours. TUBULIN and PRO-CASPASE 3 were used ad internal controls used. The graph represents the mean ± SD of three biological replicates (n = 3). Student’s t-test: ns, not significant; **p≤0.01). D) Immunofluorescence analysis to detect CLEAVED CASPASE-3 (green) in fibroblasts and MNPs treated with a 0.5 M sorbitol for 2 hours. TOM20 (red) was used to stain mitochondria, and DAPI (blue) to stain nuclei. Scale bars = 10 µm. The graph reports on the mean ± SD of the percentage of cells expressing CLEAVED CASPASE-3 in MNPs treated or not with sorbitol. n = 3 biological replicates. Student’s t-test: ns, not significant; ****p≤0.001).

We also assessed apoptosis by analyzing cleaved caspase-3 levels. In contrast to fibroblasts, MNPs showed accumulation of cleaved caspase-3 as early as 2 hours after sorbitol treatment, with a more pronounced increase at 3 hours (**Figure 5C**). Early activation of cleaved caspase-3 in MNPs was confirmed through immunofluorescence analysis and relative quantitation (**Figure 5D**).

Taken together, these findings demonstrate that neural cells are more susceptible than fibroblasts to the stress induced by sorbitol that can occur in patients with SORD deficiency. This highlights the heightened vulnerability of MNPs to damage mediated by the polyol pathway.

### Aldose reductase inhibition rescues mitochondrial alterations in SORD-deficient cells

Inhibition of AR has been shown to normalize intracellular sorbitol levels in SORD-deficient fibroblasts derived from patients^7,15^. To determine whether AR inhibition could also alleviate mitochondrial defects, we treated fibroblasts from all three patients with 100 µM of the AR inhibitor (ARi) epalrestat. Immunofluorescence analysis using the mitochondrial marker TOM20 revealed that AR inhibition fully rescued the mitochondrial phenotype in SORD-deficient fibroblasts (**Figure 6A**). Specifically, treated cells lost their characteristic fragmented mitochondrial morphology and instead displayed a complex, interconnected network resembling that of control fibroblasts (**Figure 6A**). Dimensional analysis to quantitatively evaluate these effects demonstrated that ARi treatment significantly increased mitochondrial area and perimeter (**Figure 6B** and **S5E**), indicating a restoration of mitochondrial size. In addition, mitochondrial network architecture was improved in ARi treated cells, as evidenced by increased aspect ratio and branch length (**Figure 6B** and **S5E**). The increase in aspect ratio is consistent with enhanced mitochondrial fusion^43^. Accordingly, mitochondrial circularity was reduced following AR inhibition, further supporting the rescue of normal mitochondrial network (**Figure 6B** and **S5E**).

**Figure 6.**
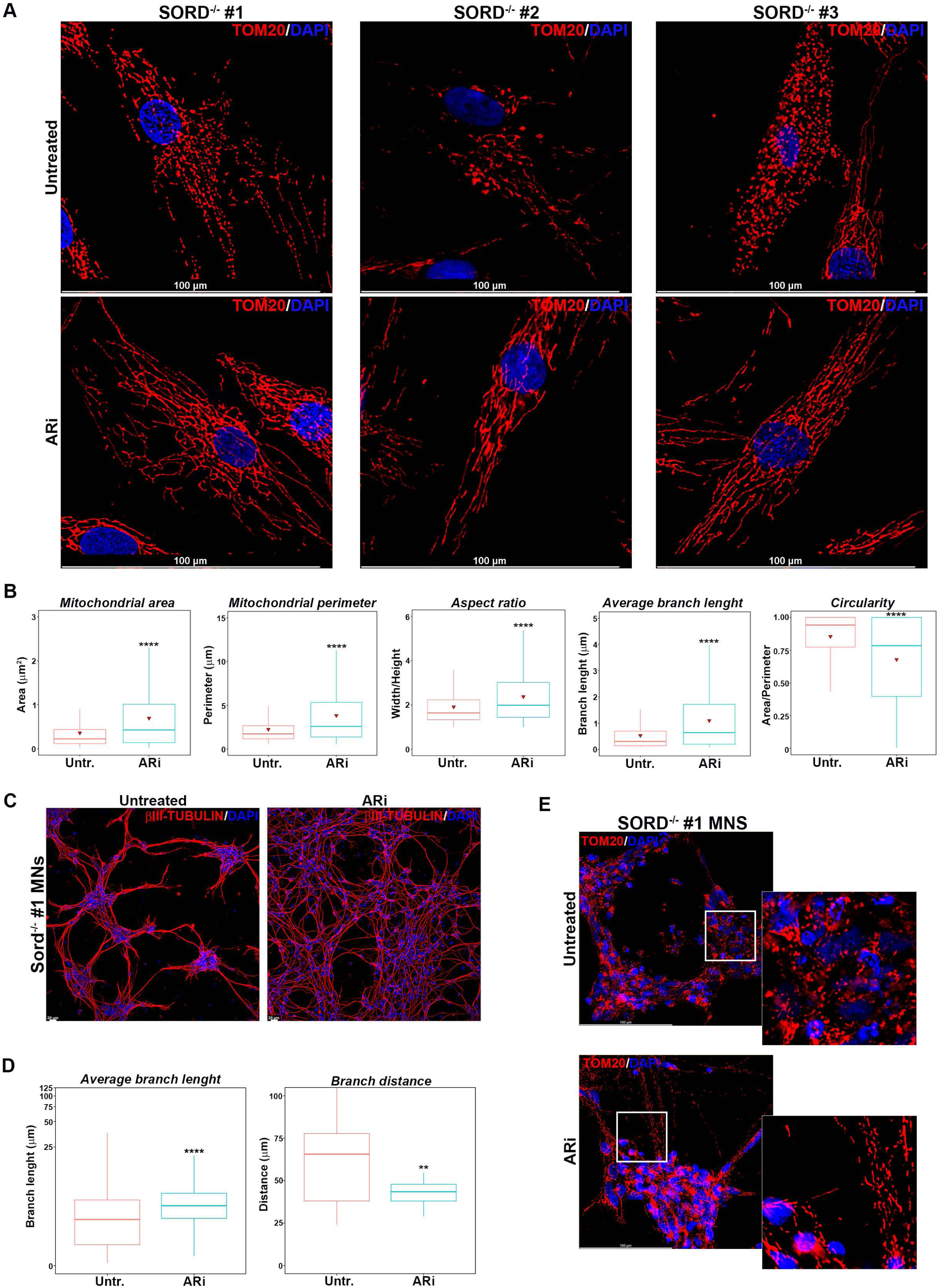
Treatment with ARi rescues the mitochondrial and cytoskeletal phenotypes observed in cells from patients with CMT2/SORD. A) Immunofluorescence analysis to evaluate mitochondrial network in SORD-deficient fibroblasts treated or not with the AR inhibitor epalrestat. Scale bars = 10 µm. B) Morphometric analysis of mitochondrial network relative to the immunofluorescence in the panel (A). Box plots represent the quantification of mitochondrial area, perimeter, aspect ratio (width/height), branch length, and circularity (area/perimeter) following treatment. The values are presented as the interquartile range (IQR, 25th–75th percentile), with the horizontal line indicating the median. Whiskers extend to the minimum and maximum values, and red triangles denote the mean for each group. Student’s t-test: ns, not significant; ****p≤0.0001). C) Immunofluorescence analysis of the neurite network of SORD-deficient MNs treated or not with ARi. βIII-TUBULIN (red) was used to stain the neuronal cytoskeleton, and DAPI (blue) to stain nuclei. Scale bars = 20 µm. D) Morphometric analysis of the immunofluorescence presented in the panel C. The neurite length and the distance between neurites were evaluated. The values are presented as the interquartile range (IQR, 25th–75th percentile), with the horizontal line indicating the median. Whiskers extend to the minimum and maximum values, and red triangles denote the mean for each group. Student’s t-test: ns, not significant; **p≤ 0.01; ****p≤0.0001). E) Immunofluorescence analysis to evaluate the mitochondrial network in SORD-deficient MNs treated or not with ARi. Scale bars = 100 µm. White square indicates the cropped region.

AR inhibition has been reported to improve nerve structure and function in diabetic neuropathy^44,45^, thus we assessed whether epalrestat could also rescue neuronal defects in SORD-deficient MNs. Notably, ARi treatment significantly improved neuronal network architecture, increasing neurite branch length and reducing branch distance in patient-derived MNs (**Figure 6C,D**).

We then investigated whether inhibiting AR could also have a positive impact on mitochondrial network in SORD-deficient MNs. Immunostaining for TOM20 followed by quantitative analysis revealed increased mitochondrial area and perimeter upon ARi treatment (**Figure 6E** and **S5F**). Moreover, AR inhibition promoted elongation of mitochondrial branches and significantly reduced mitochondrial circularity, further indicating a rescue of normal mitochondrial network (**Figure S5F**). Taken together, these findings suggest that SORD deficiency leads to defects in neuronal branching and mitochondrial network organization, accompanied by increased cell death. Furthermore, these alterations can be effectively restored by pharmacological inhibition of AR.

## Discussion

Axonal Charcot–Marie–Tooth disease (CMT) caused by mutations in the *SORD* gene is considered to be the most common autosomal recessive neuropathy. It is characterized by the accumulation of sorbitol in fibroblasts, serum, and blood of affected individuals, irrespective of sex or variant type^7,46^. Animal models, including *Sord*−/− Sprague–Dawley rats and *Drosophila*, have shown that loss of SORD enzymatic activity primarily leads to motor peripheral neuropathy. In particular, Sord-deficient flies exhibit a neurodegenerative phenotype associated with mitochondrial dysfunction, increased reactive oxygen species (ROS), and apoptosis^15^.

While these models have provided important insights into SORD deficiency, the development of a human neural model is essential to dissect disease mechanisms in detail and to enable preclinical therapeutic studies. To address this need, we generated patient-derived fibroblasts carrying *SORD* mutations and, through reprogramming and differentiation, established a human *in vitro* neural model to investigate the consequences of SORD deficiency.

We first confirmed that biallelic *SORD* mutations result in complete loss of the protein, whereas heterozygous mutations lead to reduced SORD expression. Loss of SORD function caused pronounced cytoskeletal abnormalities in patient-derived MNs, including impaired neuronal branching and altered synaptic junctions. Both fibroblasts and MNs derived from patients exhibited marked disruption of the mitochondrial network. Morphometric and functional analyses demonstrated that SORD deficiency affects mitochondrial size, ultrastructure, and respiratory capacity, leading to increased proton leak and elevated oxidative stress. Notably, electron microscopy revealed that mitochondrial defects were particularly severe in axons, while cell bodies were relatively spared, suggesting a mechanistic link to neurite degeneration.

Recently, it has been suggested that mitochondrial defects may be a common feature of axonal CMT neuropathies^33,47^. Indeed, many of the genes involved in the pathogenesis of different CMT subtypes play a role in mitochondrial biology. These genes influence processes such as mitochondrial remodelling (e.g. MFN2 and GDAP1), mitochondrial transport along peripheral axons, ATP production and oxidative stress pathways (e.g. RAB7 and GDAP1) ^17,18,48,49^. Alteration of these mechanisms prevents the axon from sustaining its high energy demand, which leads to the characteristic axonal dysfunction of many CMT2 types^50^. Our results suggest that also CMT2/SORD belongs to the group of CMT forms associated with alterations of the mitochondrial compartment. Therefore, it is important to determine whether mitochondrial defects truly contribute to the pathogenesis of the disease, or if the impairment of these functions is simply an endpoint of axonal degeneration. Further studies are needed in the case of CMT2/SORD to establish whether this neuropathy is a direct consequence of sorbitol accumulation, or if other mechanisms also contribute. Given that elevated sorbitol levels are associated with osmotic stress, toxicity, ROS production, and mitochondrial dysfunction^15^, we hypothesized that sorbitol accumulation drives the observed phenotype. Consistent with this hypothesis, exogenous sorbitol treatment in WT fibroblasts and neural cells recapitulated key features of SORD deficiency, including mitochondrial network disruption and dose-dependent mitochondrial fragmentation. Neural cells were significantly more sensitive to sorbitol than fibroblasts, with MNs undergoing apoptosis at relatively low concentrations. This heightened vulnerability may explain the neuron-specific degeneration observed in CMT patients with SORD mutations, as MNs are insulin-sensitive^51^ and take up substantial amounts of glucose, making them particularly susceptible to dysregulation of the polyol pathway.

Collectively, our findings establish a mechanistic link between SORD deficiency, sorbitol accumulation, and neural cell degeneration. SORD loss leads to increased ROS production, impaired oxidative phosphorylation (OXPHOS), indicating mitochondrial dysfunction. Importantly, mitochondrial abnormalities were observed both in SORD-deficient cells and following sorbitol exposure, and both conditions were associated with increased neural cell death. The connection between SORD deficiency and sorbitol accumulation is further supported by the accumulation of AR observed in both contexts. In the absence of SORD, sorbitol cannot be converted to fructose, leading to its accumulation and the establishment of a feed-forward loop whereby increased AR activity further leads to sorbitol accumulation (**Figure 7**). This self-reinforcing cycle likely contributes to cellular damage. Moreover, elevated AR activity may impair antioxidant defences through NADPH depletion, thereby limiting glutathione (GSH) regeneration and exacerbating oxidative stress^10,52^ (**Figure 7**). These observations suggest the possibility of testing mitochondria-targeted antioxidants^53,54^ for CMT/SORD disease.

**Figure 7.**
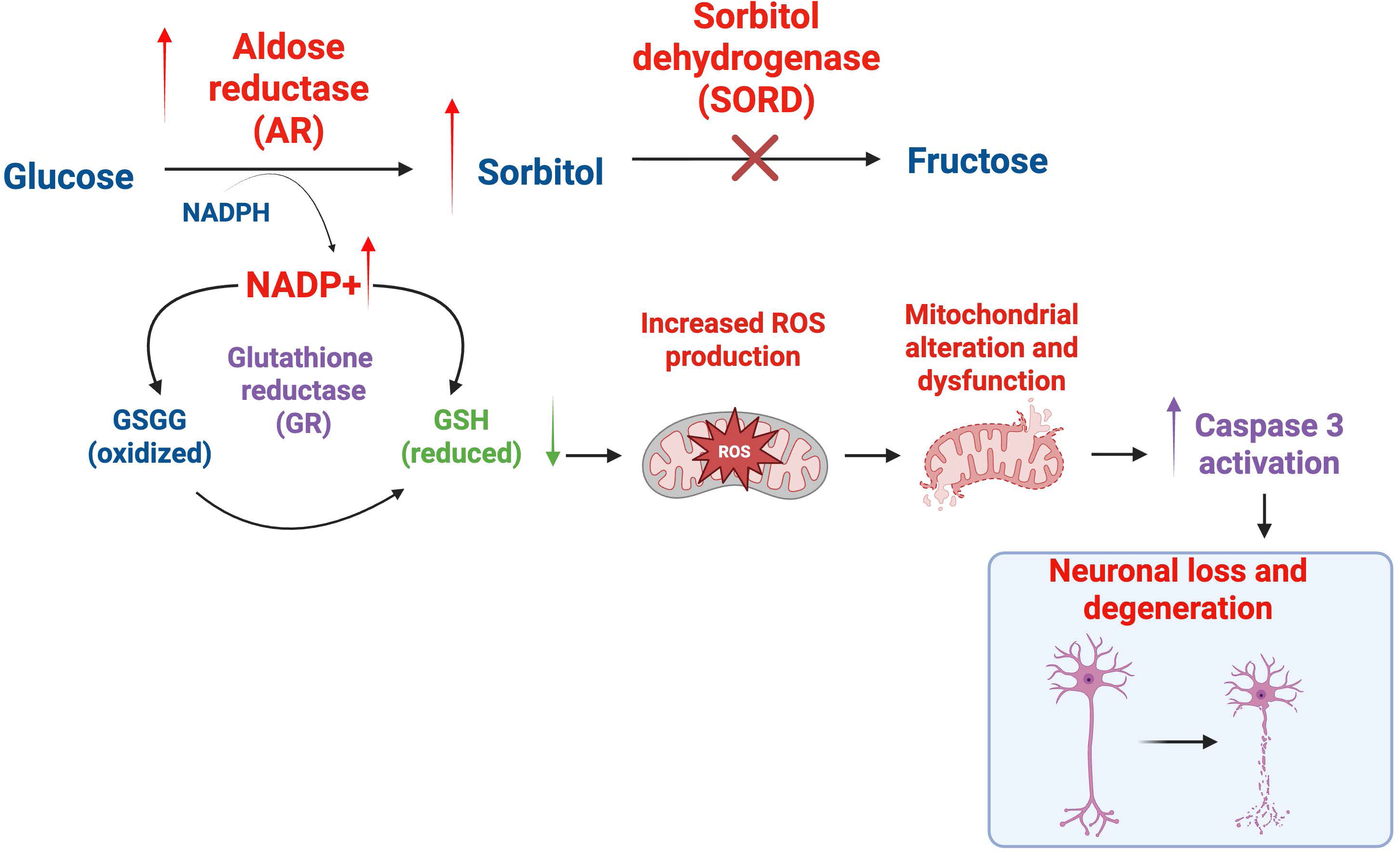
Proposed model illustrating the effects of SORD deficiency in MNs. In the absence of SORD, sorbitol cannot be converted to fructose, leading to its accumulation and the establishment of a feed-forward loop in which increased AR activity further promotes sorbitol production. This cycle likely contributes to cell damage. Moreover, increased AR activity may impair antioxidant defences by consuming NADPH, thereby limiting glutathione (GSH) regeneration and exacerbating oxidative stress. Coloured arrows indicate dysregulated changes in pathway components resulting from SORD deficiency (red, increased; green, decreased), while the violet arrow denotes aberrant cell death associated with SORD deficiency.

Based on our findings and previous studies, inhibition of AR activity, and consequent reduction of sorbitol accumulation, represents a promising therapeutic strategy for SORD deficiency. Here, we demonstrated that the treatment with the AR inhibitor epalrestat restored neurite integrity and mitochondrial network organization in SORD-deficient MNs. Furthermore, *Zhu and colleagues*^15^ demonstrated that the AR inhibitor AT-007 reduced sorbitol levels by approximately 50% in MNs and 35% in the *Drosophila* brain, mitigating sorbitol-induced toxicity. Notably, AT-007 (govorestat) is currently under clinical investigation (NCT05397665) to evaluate its effects on sorbitol reduction and clinical outcomes in patients with SORD deficiency.

It is also worth noting that our results provide important insights into the functional mechanisms underlying diabetic neuropathy and its therapeutic treatment. Indeed, the key features that we identified in CMT2/SORD, including the excessive conversion of glucose to sorbitol and the subsequent overexpression of the AR enzyme, resulting in osmotic stress and oxidative damage, also represent fundamental hallmarks of diabetic neuropathy^55–57^. Furthermore, as observed in CMT2/SORD neuropathy, individuals who express high levels of AR often develop diabetic neuropathy at an early stage^58,59^. Together with findings supporting the inhibition of AR to prevent excessive intracellular sorbitol accumulation, this evidence suggests that targeting this enzyme could represent a powerful therapeutic strategy for neuropathies characterized by dysfunction of the polyol pathway^60–62^. This is particularly relevant given that, while AR inhibitors (e.g., epalrestat) are used to treat diabetic neuropathy in some countries, they have not yet been widely recognized as a definitive therapeutic approach worldwide (https://biologyinsights.com/epalrestat-uses-side-effects-and-availability/).

Overall, our findings support the therapeutic potential of AR inhibitors as a targeted pharmacological strategy for polyol pathway-related neuropathies. They also highlight the feasibility of developing therapies that target shared pathogenic mechanisms in inherited neuropathies, including mitochondrial dysfunction and oxidative stress, which could ultimately benefit a broader population of patients with Charcot–Marie–Tooth (CMT) disease.

## Data availability

The data that support the findings of this study are available from the corresponding author, upon reasonable request.

## Acknowledgements

The authors would like to thank: the “BIOmedical iMAGIng fAcility” of DMMBM (Naples, Italy); Ludovica D’Auria and Caterina Missero of the Advanced Light Microscopy Facility of CEINGE (Naples, Italy) for help with imaging; Danilo Swann Matassa for critical discussion of the experiments.

## Funding

This research was supported by Ministero dell’Università e della Ricerca (MUR) PRIN 2022 #2022P7R5CJ to SP and PRIN/PNRR2022 #P2022KBAT7 to FM; Italian Association for Cancer Research (AIRC), Investigator Grant #24976 and by the Ministero dell’Università e della Ricerca (MUR), PRIN/PNRR2022 #P2022F3YRF to PM. ST is supported by the Ministry of University and Research (MUR), National Recovery and Resilience Plan (NRRP), project MNESYS (PE0000006) - A multiscale integrated approach to the study of the nervous system in health and disease (DN. 1553 11.10.2022)

## Author contributions

**Giuseppina Divisato:** experimental design, data collection, analysis and interpretation, manuscript writing and editing. **Stefano Tozza:** data collection, analysis and interpretation, manuscript writing and editing. **Elena Polishchuk, Maria Chiara Zizolfi, Emilia Giannino, Fiorella Marsella, Daniela Di Girolamo, Ciro Menale, Lucia Perone, Paolo Gianfico:** data collection, validation and analysis. **Giovanni Cuda, Cecilia Bucci, Paolo Maiuri, Roman Polishchuk:** experimental design, data interpretation, manuscript editing. **Fiore Manganelli:** experimental design, data interpretation, resources, manuscript writing and editing. **Silvia Parisi:** conceptualization and experimental design, data collection, analysis and interpretation, resources, project administration, manuscript writing and editing.

## Competing interests

The authors declare no competing interests

## Ethics declarations

The experiments were performed according to the approved study protocols by the Ethics Committee of University of Naples Federico II (334/21).

## Supplementary material

Supplementary material is available in “Supplementary material” PDF file.

## Notes

### Competing Interest Statement

The authors have declared no competing interest.

