## Supplementary Figures and legends for "Motor Neuron Dysfunction in SORD Deficiency: Implications for Therapeutic Development in Peripheral Neuropathies"

#### Supplementary Figure 1

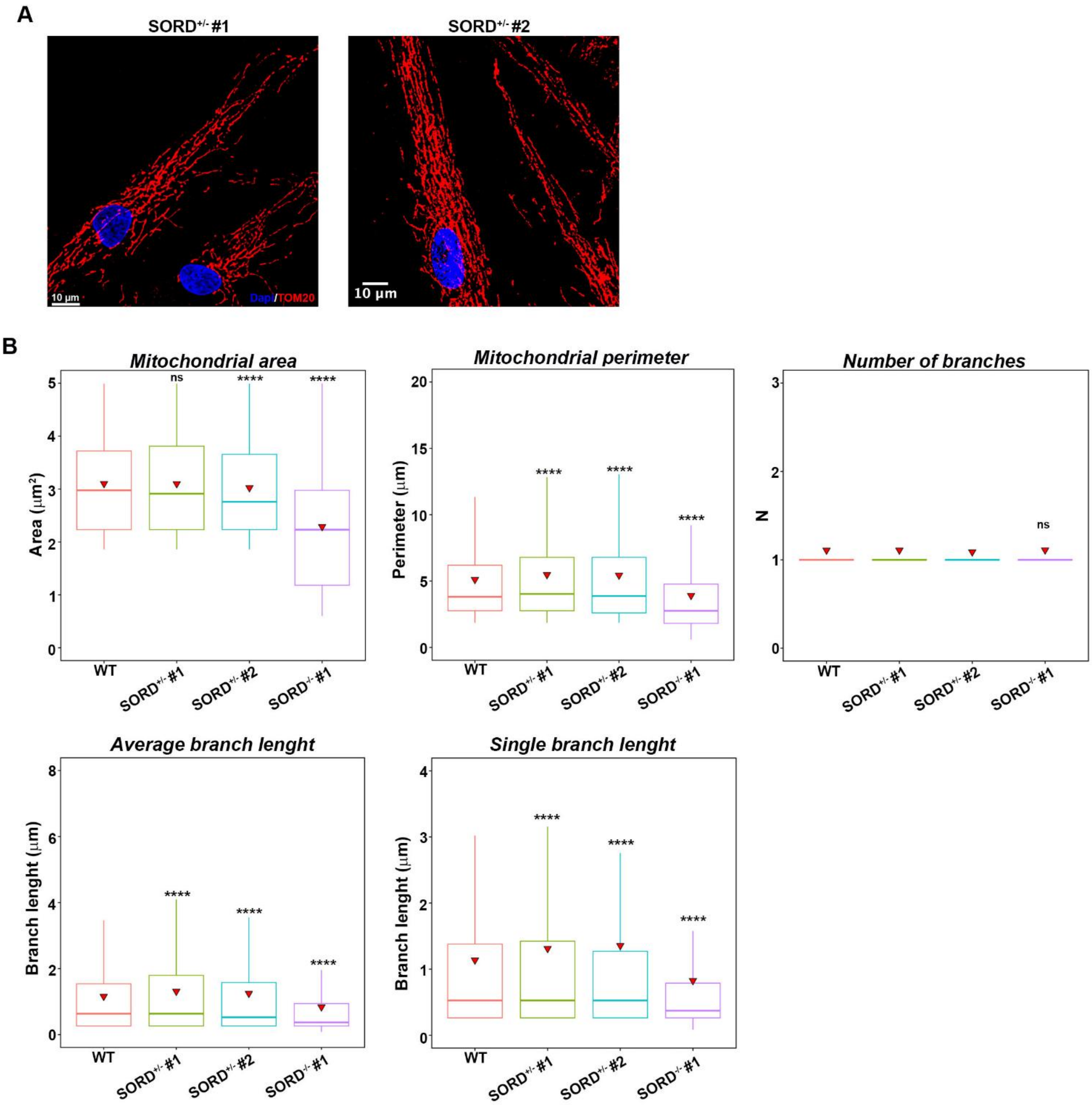


**Figure S1. Analysis of mitochondria in WT and SORD-deficient cells.**

A) Immunofluorescence analysis relative to the Figure 1D to evaluate the mitochondrial network in fibroblasts of *SORD*-mutated gene carriers. The mitochondria were stained with TOM20 antibody (red) and the nuclei with DAPI (blue). Scale bars = 10 µm.

B) Morphometric analysis of mitochondrial compartment in fibroblasts stained with Mitotracker™ Green FM (see **Supplementary Movies**). Box plots report on the evaluation of mitochondrial dimesions (area and perimeter) along and single branch length. The values represent the interquartile range (IQR, 25th–75th percentile), with the horizontal line indicating the median. Whiskers extend to the minimum and maximum values, and red triangles denote the mean for each group. Student's t-test: ns, not significant; ****$p \leq 0.0001$.

### Supplementary Figure 2

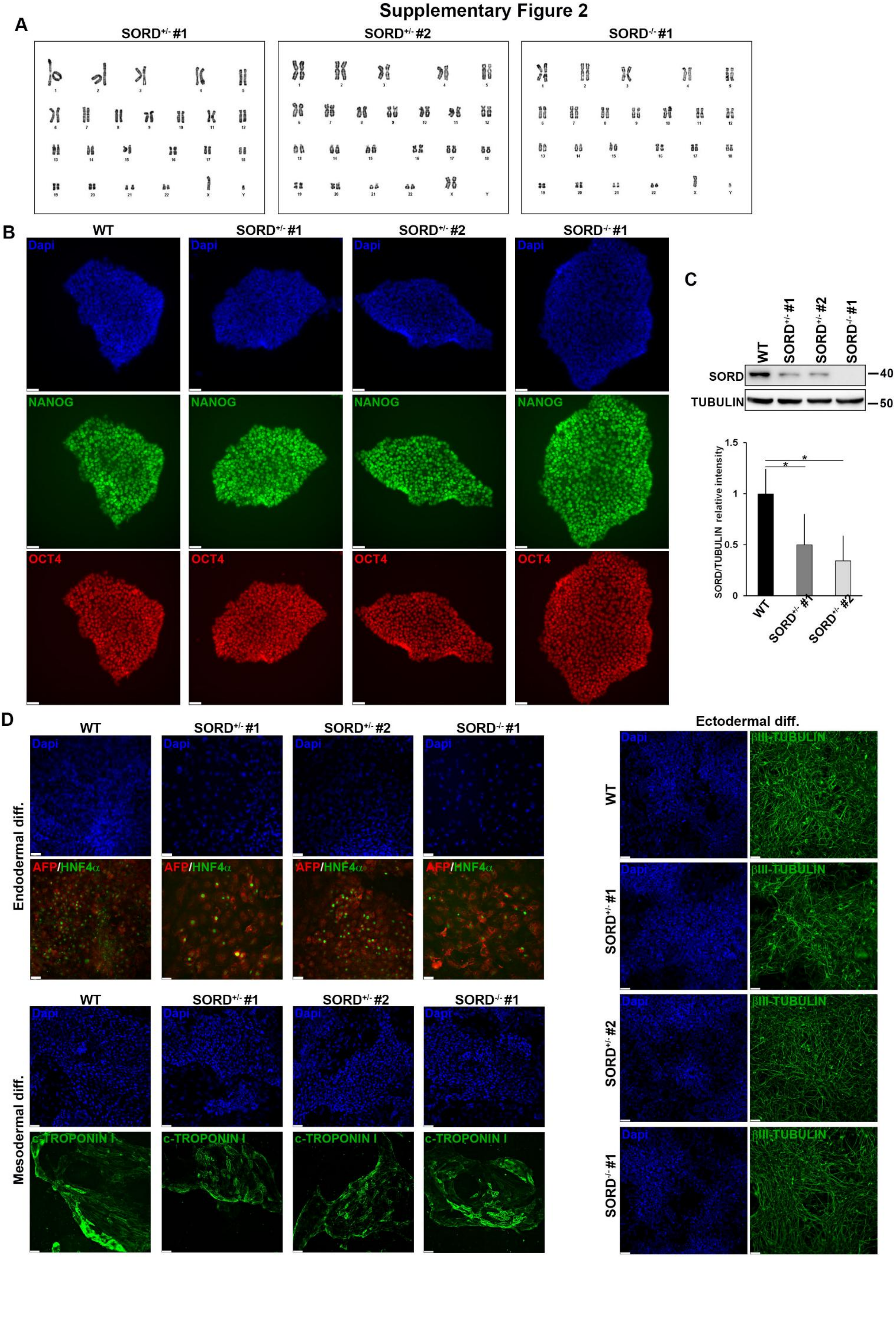

**Figure S2. Characterisation of iPSCs generated from fibroblasts of WT and SORD-mutated individuals.**

A) Karyotype analysis of the iPSCs generated from fibroblasts of the two unaffected carriers and the CMT2/SORD patient to evaluate chromosomes number and structure.

B) Immunofluorescence analysis of the stemness markers NANOG (green) and OCT4 (red) in WT, carrier ($SORD^{+/-}$ #1, $SORD^{+/-}$ #2) and patient ($SORD^{-/-}$ #1). DAPI (blue) was used to stain the nuclei. Scale bars: 50 µm.

C) Western blot analysis and the relative quantification to evaluate the expression of SORD in the indicated iPSCs. The graph reports on the mean ± SD of SORD relative to total TUBULIN signal intensity of three biological replicates (n=3). Student's t-test: ns, not significant; *$p\leq0.05$, **$p\leq0.01$).

D) Immunofluorescence analysis  to assess the differentiation of the iPSCs into the three germ layer derivatives. AFP (red) and HNF4a (green) were used as endodermal markers; βIII-TUBULIN (green) was used as neuronal marker and CARDIAC TROPONIN (c-TROPONIN) was used as mesodermal marker. DAPI was used to stain nuclei. Scale bars: 50 µm.

Supplementary Figure 3

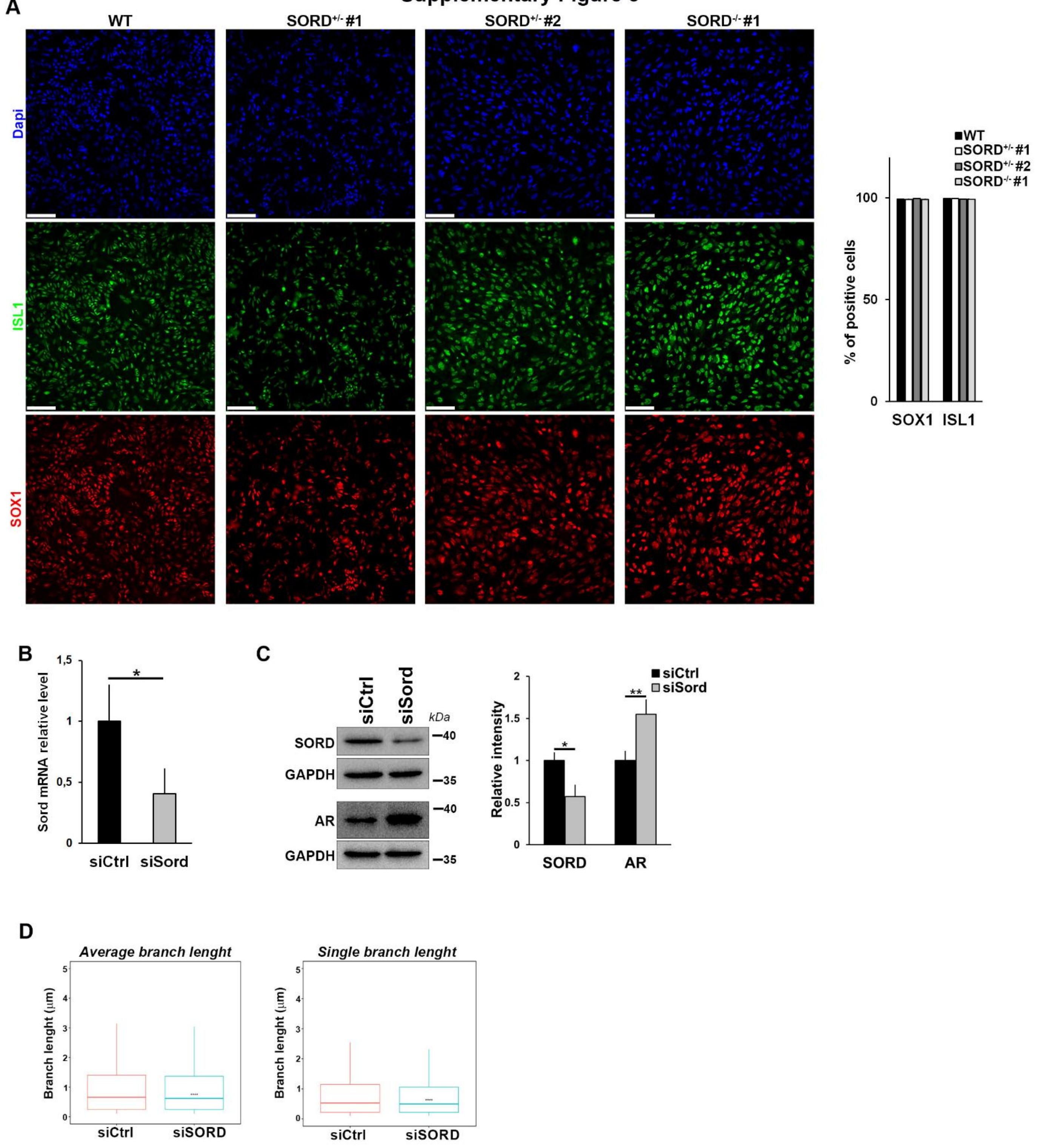


**Figure S3. Characterisation of human MNPs and MNs obtained from WT and *SORD*-mutated iPSCs.**

A) Immunofluorescence analysis and relative quantification to evaluate the expression of MNP markers ISL1 (green) and SOX1 (red) in MNPs obtained from the indicated iPSC lines. DAPI was used to stain nuclei. Scale bar=100 µm. The graph shows the percentage of cells expressing SOX1 and ISL1 markers on total cells. Data are presented as mean ± SD (n = 3 biological replicates). Student's t-test: not significant.

B) Quantitative qPCR to evaluate the levels of *SORD* mRNA in MNPs transfected with a control or a *SORD*-specific siRNA after 72 hours from transfection. The graph represents the mean ± SD normalized to Gapdh and relative to the siCtrl sample in n=3 biological replicates. Student's t-test: *$p \leq 0.05$).

C) Western blot analysis and relative quantification to evaluate the level of SORD and AR proteins in MNPs transfected with a control or a *SORD*-specific siRNA after 96 hours from transfection. GAPDH levels were used as loading control. The graph represents the mean ± SD relative to the siCtrl sample (n=3 biological replicates). Student's t-test: *$p<0.05$; **$p<0.01$.

D) Morphometric analysis of the mitochondrial network in MNPs transfected with siCtrl and si*SORD*, relative to the immunofluorescence in **Figure 2C**, after 96 hours from transfection. The values are presented as the interquartile range (IQR, 25th–75th percentile), with the horizontal line indicating the median. Whiskers extend to the minimum and maximum values, and red triangles denote the mean for each group. Student's t-test: ****$p \leq 0.0001$.

### Supplementary Figure 4

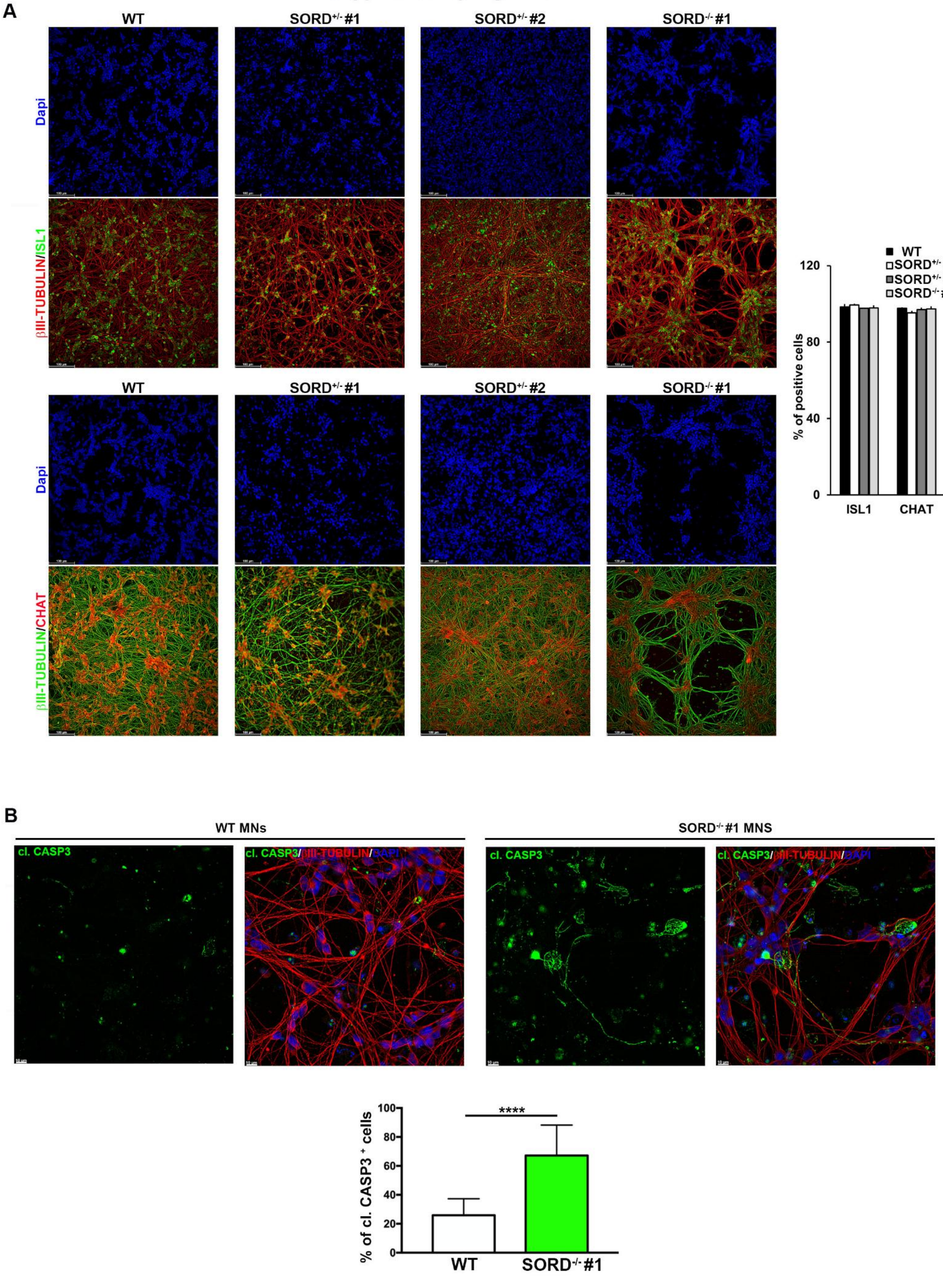

**Figure S4. SORD-deficient MNs show cytoskeletal abnormalities and increased cell death.**

A) Immunofluorescence analysis to evaluate cytoskeleton in MNs obtained from WT, carrier ($SORD^{+/-}$ #1, $SORD^{+/-}$ #2) and patient ($SORD^{-/-}$ #1) iPSCs. βIII-TUBULIN (red in upper panel and green in lower panel) was used to stain neuronal cytoskeleton, while ISL1 (green, upper panel) and CHAT (red, lower panel) were used as markers of differentiated MNs. DAPI (blue) was used to stain nuclei. Scale bars=100 µm. The graph shows the percentage of cells expressing the MN markers ISL1 and CHAT for each condition. Data are presented as mean ± SD (n = 3 biological replicates). Student's t-test: not significant.

B) Immunofluorescence analysis to assess the presence of CLEAVED CASPASE-3 (green) in WT and SORD-deficient MNs. βIII-tubulin (red) was used to stain the neuronal cytoskeleton, and DAPI (blue) was used to stain nuclei. Scale bars = 10 µm. The graph reports the percentage of cells expressing CLEAVED CASPASE-3 on total cells. Data are presented as mean ± SD (n = 3 biological replicates). Student's t-test: ****$p \leq 0.001$.

#### Supplementary Figure 5

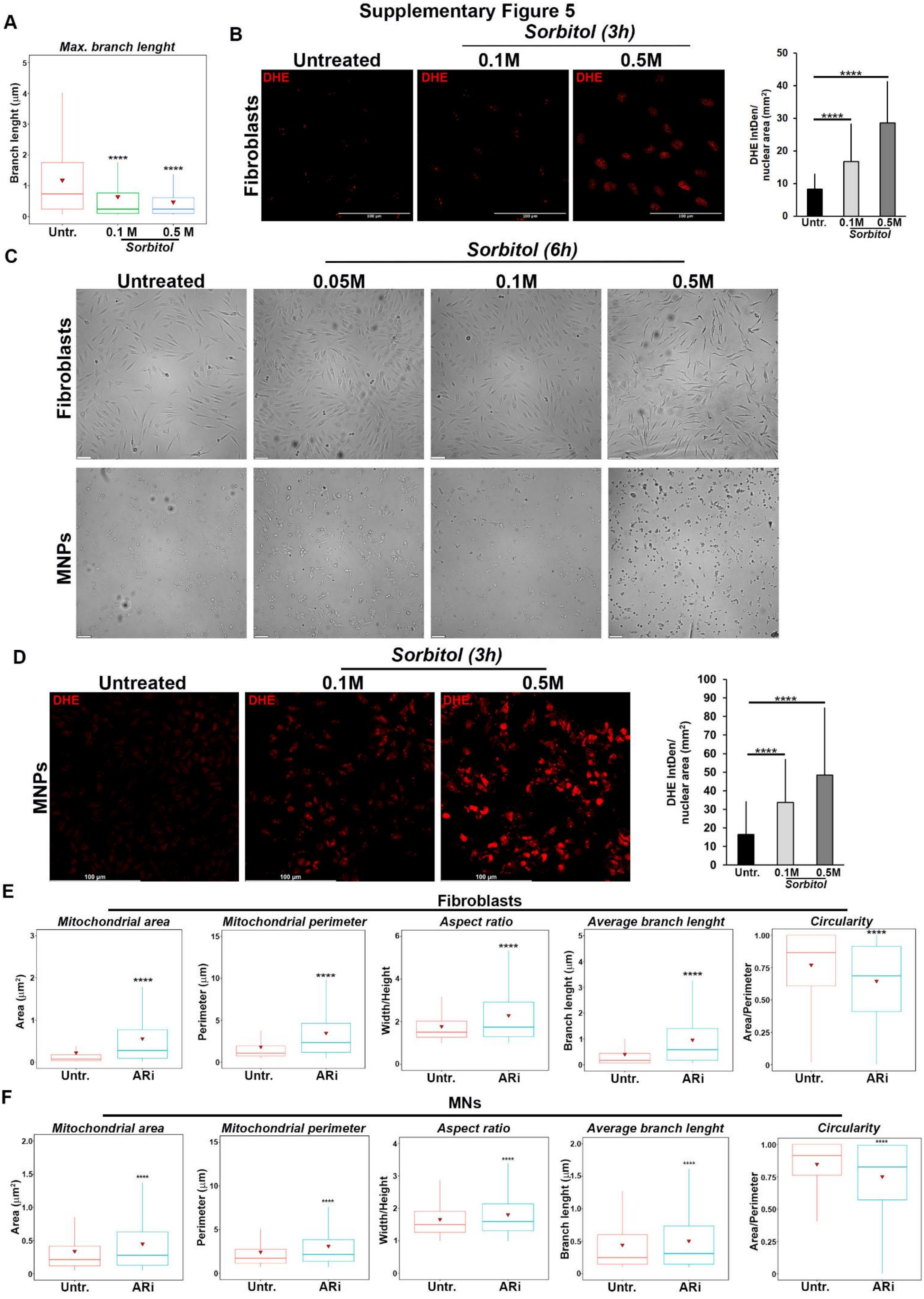

**Figure S5. Sorbitol accumulation induces an early, dose-dependent increase of ROS in neural cells.**

A) Morphometric analysis of mitochondrial branch length of fibroblasts treated with the indicated concentrations of sorbitol and relative to the immunofluorescence reported in Figure 4A. Student's t-test: ****p≤0.001.

B) DHE staining of fibroblasts treated with the indicated concentrations of sorbitol for three hours. Scale bars = 100 μm. The graph reports the DHE signal intensity (integrated density/nuclear area, $mm^2$) in treated fibroblasts compared to untreated cells. Data are presented as mean ± SEM (n = 2 biological replicates). Student's t-test: ****p≤0.001.

C) Phase-contrast images of WT fibroblasts and MNPs treated with the indicated concentrations of sorbitol for 6 hours. Scale bars = 100 μm.

D) DHE staining of MNPs treated with the indicated concentrations of sorbitol for three hours. The graph shows the quantitation of DHE signal intensity (integrated density/nuclear area, $mm^2$). Data are presented as mean ± SEM (n = 2 biological replicates). Student's t-test: ****p≤0.001.

E) Morphometric analysis representing mitochondria parameter of both $SORD^{-/-}$ #2 and 3 fibroblasts treated or not with ARi and relative to the immunofluorescence reported in **Figure 6A**. Box plots report the evaluation of mitochondrial area, perimeter, aspect ratio (width/height), branch length and mitochondrial circularity (area/perimeter). The values are presented as the interquartile range (IQR, 25th–75th percentile), with the horizontal line indicating the median. Whiskers extend to the minimum and maximum values, and red triangles denote the mean for each group. Student's t-test: ****p≤0.001.

F) Morphometric analysis of mitochondria in $SORD^{-/-}$ #1 MNs treated or not with ARi and relative to the immunofluorescence reported in **Figure 6E**. Box plots show changes in mitochondrial parameters, including area, perimeter, aspect ratio (width/height), average branch length, and circularity (area/perimeter). The values are presented as the interquartile range (IQR, 25th–75th percentile), with the horizontal line indicating the median. Whiskers extend to the minimum and maximum values, and red triangles denote the mean for each group. Student's t-test: ****p≤0.001.
